# Exposure to p,p’-dichlorodiphenyldichloroethylene (p,p’-DDE) Perturbs Cell State and Stem Cell Markers and Induces an Epidermal Phenotype in Human Breast Cells

**DOI:** 10.64898/2026.10.13.758898

**Authors:** Jenna L. Miller, Anagha Tapaswi, Jesse W. Wotring, Katelyn M. Polemi, Penghui Yang, Xiang Zhou, Laurie K. Svoboda, Jonathan Z. Sexton, Justin A. Colacino

## Abstract

Breast cancer is the second leading cause of cancer deaths in U.S. women. Endocrine disruptors, such as the DDT metabolite DDE, are environmental factors linked to breast cancer risk. Emerging evidence shows that during carcinogenesis, cells reacquire stem cell traits and exhibit phenotypic plasticity. In these phenotypically plastic states, breast cells transition between phenotypes, for example, from epithelial to mesenchymal (EMT) phenotypes, or from luminal to basal. Discovering how environmental factors alter cell phenotype may be a critical step in preventing and treating breast cancer. We applied a novel high-content imaging method for the evaluation of expression of stemness and cell state markers (ALDH1A3, CD44, CD24, KRT8, and KRT14) and evaluated these markers after introducing environmentally relevant concentrations of DDE (25nM to 25µM) to non-tumorigenic MCF10A breast epithelial cells grown in 3D anchorage independent conditions. After identifying surprisingly profound cell state alterations, including a loss of KRT8 and KRT14 expression, by 25 µM DDE exposure, we used transcriptomic profiling along with pharmaceutical and genetic knockdown approaches to characterize the molecular mechanisms by which DDE exposure modulates breast epithelial cell state. Keratinization and formation of the cornified envelope were biological pathways upregulated with a maximum fold change of 360.47. We identify *DLX3* as a transcription factor which is inducible by DDE exposure and is also highly upregulated in human breast cancers, reflecting a novel mechanism by which environmental exposures may promote cellular plasticity and breast carcinogenesis.

## Introduction

Breast cancer is the most common cancer diagnosed among women in the U.S. and about 42,140 women are estimated to die from breast cancer in 2026 (1). While some known risk factors for breast cancer include inherited genetics, age, sex, family history, smoking, drinking, and obesity, there are many environmental exposures that contribute to breast cancer development (2) with poorly understood mechanisms of action.

Environmental exposures can act as genotoxic agents, alter mammary gland development or hormone responsiveness, and promote breast cancer (3). p,p’-dichlorodiphenyldichloroethylene (DDE) is a metabolite of dichlorodiphenyltrichloroethane (DDT), a persistent insecticide classified as an IARC group 2 carcinogen (4). Epidemiological studies have found that high levels of DDT exposure before puberty greatly increases breast cancer risk in women (5). We and others find that DDE is readily detectable in US women with biological concentrations variable by demographic factors such as age and race, and maximum blood lipid p,p’-DDE concentrations exceeding 30 µM (6–10). DDE concentrations in breast adipose tissue are around 2.3 µM in breast cancer patients (11), with levels estimated to be higher in breast cancer patients versus controls (12). Thus, there is a need to understand mechanisms by which DDT and its metabolites promote the development of breast cancer, particularly when exposures are occurring during windows of susceptibility.

One of these mechanisms could be by impacting cellular plasticity (13,14). Building evidence shows that during carcinogenesis, cells reacquire stem cell-like traits and exhibit phenotypic plasticity, a Hallmark of Cancer (15). Stemness and plasticity is reflected by transitions between epithelial and mesenchymal states or luminal and basal states, with cells exhibiting “hybrid” properties or the simultaneous expression of multiple cellular states (16–18). For example, normal breast and breast cancer stem cells have been characterized by their expression of markers including aldehyde dehydrogenase (ALDH, including ALDH1A3), CD44, and CD24, which reflect epithelial stem-like (ALDH+), mesenchymal stem-like (CD44+/CD24-), and hybrid epithelial/mesenchymal (ALDH+/CD44+/CD24-) states (16,19). In breast cancer, the ALDH+ and CD44+/CD24-cancer stem cell populations are typically non-overlapping (19), whereas a significant population of normal breast cells have ALDH+/CD44+/CD24-expression and a “hybrid” epithelial/mesenchymal phenotype (16). Related to luminal/basal states, the luminal epithelium of a mammary gland expresses the keratin 8 (KRT8) and the myoepithelial/basal layer expresses keratin 14 (KRT14). Basoluminal hybrid cells co-express these markers, indicating loss of cellular identity (20). Aggressive breast cancers have expression patterns which mirror stem cells or “hybrid” cells (16,18). Understanding how exposures, like DDE, dysregulate cellular plasticity and stemness would provide important insights into mechanisms of breast carcinogenesis.

Here, we applied a three-dimensional *in vitro* approach using non-tumorigenic breast cells to characterize the impacts of DDE exposure on stemness and plasticity markers. We developed a high-content imaging method for the evaluation of stemness and plasticity marker protein expression, testing concentration-dependent effects of DDE. After surprisingly identifying profound cell state alterations induced by DDE exposure, we used transcriptomic profiling along with pharmaceutical and genetic knockdown approaches to characterize the molecular mechanisms by which DDE exposure modulates breast epithelial cell state. We identify *DLX3* as a transcription factor which is inducible by DDE exposure and is also highly upregulated in human breast cancers, reflecting a novel mechanism by which environmental exposures may promote cellular plasticity and breast carcinogenesis.

## Materials and Methods

### Cell Culturing and Mammosphere Formation

Non-tumorigenic mammary epithelial cells, MCF10A (ATCC, CRL-10317), were maintained in 2D as previously described (10). For 3D culture as “mammospheres”, cells were resuspended in MammoCult media (STEMCELL, NC9021910) with or without DDE. DDE (Chem Service, N-10875-100MG) concentrations were as follows: 25 µM, 2.5 µM, 0.25 µM, and 0.025 µM DDE. 0.5% dimethyl sulfoxide (DMSO) (Sigma Aldrich, D2650). Specific details about the mammosphere cultures can be found in the Supplemental Methods.

### Mammosphere Staining and Imaging

To quantify mammosphere number and size in an unbiased manner, mammospheres were stained with Hoechst 33342 (Fisher, H3570) and stained plates were imaged using an Invitrogen EVOS M7000. Details about staining and image acquisition can be found in the Supplemental Methods.

### Stem Cell and Cell State Marker Immunostaining and Imaging

One replicate was collected from the 6-well plate after seven days of incubation. The collected spheres were centrifuged at 110 g for 10 minutes, washed one time with 1X PBS and resuspended in DDE mammocult media. The spheres were plated on two collagen-coated adherent black clear-bottom 96-well plates (Sigma Aldrich, 122-20; Corning, 07-200-588) and incubated for 24 hours to ensure attachment to the bottom of the plate. Immunostaining of mammospheres was performed in three biological rounds with three technical replicates per condition. Two staining panels were developed – one assessing luminal and basal state and one assessing epithelial/mesenchymal stem cell markers. For the luminal/basal “keratin” panel, mammospheres were stained with an anti-KRT8 antibody (Abcam, ab192467), an anti-KRT14 antibody (Abcam, ab192056), and the DNA stain Hoechst 33342. For the epithelial/mesenchymal “stemness” panel, mammospheres were stained with and anti-CD44 antibody (Biolegend, 103016), an anti-CD24 antibody (Santa Cruz, sc-19585 AF647), an anti-ALDH1A3 antibody (Abcam, ab129815), and the DNA stain Hoechst 33342. Three technical replicates per condition were set up for each staining panel. A Yokogawa CV8000 microscope was used for automated plate imaging with a ×20/1.0NA water immersion objective lens, 50-µm pinhole. Laser power for each channel was adjusted to ensure optimal signal-to-noise ratios and laser-based autofocus was performed at each field with 16 fields per well captured. Specific details about the staining protocol are shown in the Supplemental Methods.

### High Content Imaging Analysis

Raw images were exported in 8-bit TIFF format. After imaging the Hoechst-stained mammospheres on an M7000 microscope and immunostained mammospheres on the Yokogawa CV8000 microscope, the images were processed through the open-source software Cell Profiler 4.2.1 (21) on a Windows-based workstation with dual Intel Xeon Platinum 8173 M processors and 192 GB of RAM. We created a CellProfiler pipeline that applied illumination correction (only for M7000 images), flagged out overly-saturated images, and identified objects of interest before measuring their characteristics. Measurements such as object size and shape, object intensity, object intensity distribution, colocalization, and texture were recorded.

### High Content Imaging Statistical Analysis

Data output from CellProfiler were uploaded to R version 4.4.2. For mammosphere size and number quantification, generalized linear models were used to measure effects of DDE, controlling for experimental round, on mammosphere size and mammosphere area. For analysis of the cell state and stemness marker expression data, across the three rounds and three replicates, the mean intensity of KRT8, KRT14, CD44, CD24, and ALDH1A3 across all individual cells in the mammospheres of each condition were calculated. Using the cytominer package v0.2.2 (22) in R, median absolute deviation normalization was performed on mean intensity data with the robustize function and transformation was performed with the generalized log operation. Intensity data was then averaged across cells per technical replicate. Generalized linear models were run with normalized and transformed data averaged across each replicate to estimate the intensity units for KRT8, KRT14, CD44, CD24, and ALDH1A3 across four concentrations of DDE compared to control and across three experimental rounds.

Cell-type proportions were calculated after robust normalization (median/median absolute deviation) to the control population where cells were then classified as high or low for each marker relative to the control median. For the “keratin” panel, there are four possible classifications: “KRT8 high/KRT14 high”, “KRT8 high/KRT14 low”, “KRT8 low/KRT14 high”, and “KRT8 low/KRT14 low”. For the “stemness” panel, there are eight possible classifications: “ALDH high/CD44 high/CD24 high”, “ALDH high/CD44 high/CD24 low”, “ALDH high/CD44 low/CD24 high”, “ALDH high/CD44 low/CD24 low” “ALDH low/CD44 high/CD24 high”, “ALDH low/CD44 high/CD24 low”, “ALDH low/CD44 low/CD24 high”, and “ALDH low/CD44 low/CD24 low”.

To model the cell-type compositions and variability across exposures, we used the sccomp R package v2.1.27 (23). A mixed-effects model was fitted using full Bayesian (Hamiltonian Monte Carlo) inference with cell-type proportions modeled as a function of DDE concentration and biological round included as a random effect. Both the composition and variability were modeled as functions of DDE concentration. The function sscomp_test was used to test for statistical significance.

### RNA-Sequencing Analysis of DDE Effects in Mammospheres

To assess the effects of DDE exposure on mammosphere gene expression, we used the SeqWell plexWell plate-based preparation kit, following our previously defined protocol (10). Detailed methods for the library preparation and sequencing can be found in the Supplemental Methods.

### Differential Gene Expression Analysis

Per sample, per gene UMI counts were loaded into R and analyzed via the package edgeR v4.4.2 (24). A metadata file containing information about chemical type and concentrations was matched to their respective samples. A design matrix was created with the “concentration” variable included to test for differences between concentration and control. Genes with low counts were filtered out with the filterByExpr function and dispersion estimates were calculated. Next, a quasi-likelihood negative binomial generalized linear model was fit for each gene and a quasi-likelihood F-test was performed to compare differentially expressed genes between exposed and control samples as well as between mixtures and their individual components.

Volcano plots were generated using log fold change and the false discovery rate p-value (FDR) to illustrate the top-most significantly differentially expressed genes between DDE and control.

### Pathway Enrichment and Benchmark Concentration Analysis

BMDExpress3 was utilized to conduct pathway enrichment analysis based on differentially expressed genes attributed to chemical exposure. Text files were generated in R that contained sample name, concentration, gene ensemble identification numbers, and gene counts per million (CPMs). A one-way ANOVA with a p-value cutoff of < 0.05 when computed for multiple testing corrections and a fold filter cutoff of 2.0 was run to filter out non-significant changes in gene expression based on concentration. A benchmark concentration analysis was performed using ToxicR Laplace model averaging. Genes were determined to have a benchmark concentration response if gene expression changed by one standard deviation relative to control. From there, REACTOME (25) pathway analysis was conducted on filtered genes. Pathways with a Fisher’s exact two-tailed test < 0.05 were considered significantly enriched pathways.

### Cardamonin Cell Viability and Assaying β-catenin Expression

To test the hypothesis that cell state transitions caused by DDE exposure were driven by induction of Wnt activity, as we have seen previously (26), we next assessed whether the known Wnt inhibitor cardamonin (27) could inhibit these transitions. To first assess cardamonin’s concentration-dependent effects on cytotoxicity and Wnt activity, MCF10A cells in a 384 well plate were exposed to a range of cardamonin concentrations and assayed at 48 and 96 hours. Cardamonin (Sigma Aldrich, 681672-2MG) was dissolved in DMSO and the following concentrations were prepared in MCF10A media: 30, 15, 7.5, 3.75, 1.88, 0.94, 0.47, 0.23, 0.12, and 0.06 µM. 0.5% DMSO in MCF10A media was used for the vehicle control. Cells were fixed and permeabilized prior to staining with an anti β-catenin primary antibody (Thermo, 71-2700) and the nuclear stain Hoechst. Plates were imaged on a Thermo Scientific CellInsight CX5 microscope using HCS Studio: Cellomics Scan v6.6.1 at 20x objective with 9 fields per well. Culture and staining protocol details can be found in the Supplemental Methods.

### DDE and Cardamonin Mammosphere Co-exposure

The following concentrations were prepared in mammocult media: Vehicle control with 0.5% DMSO, 1 µM CHIR99021 (Cayman Chemical, #13122) which is a positive control for Wnt activation (28), 25 µM DDE, 2.5 µM DDE, 3.75 µM cardamonin, 0.94 µM cardamonin, 25 µM DDE + 3.75 µM cardamonin, 2.5 µM DDE + 3.75 µM cardamonin, 25 µM DDE + 0.94 µM cardamonin, and 2.5 µM DDE + 0.94 µM cardamonin. As indicated before, mammospheres were first seeded with 200,000 cells per well in low-adherent 6-well plates with four replicates of each condition. The mammospheres incubated for seven days with 1 mL of DDE or control media being added on the third day.

### Drug-seq cDNA and Library Preparation

During the course of the studies in this manuscript, the SeqWell RNA-seq kit used for the first round of RNA-seq was discontinued. Thus, RNA-seq measures of the effects of DDE, cardamonin, and CHIR99021 were conducted using another similar plate-based method, the MERCURIUS Drug-Seq kit (Alithea Genomics). Details on library preparation and sequencing methods for Drug-Seq can be found in the Supplemental Methods. Libraries were sequenced on an Aviti24 (Element Biosciences) Sequencer using paired end reads with offset read lengths of 29 for read 1 (sample barcode plus unique molecular identifier) and 100 for read two (gene fragment). Raw sequencing reads were simultaneously demultiplexed, aligned to a splice junction aware build of the human genome, and counted per sample using STAR in the STARsolo mode. To exclude PCR duplicates, the UMI counts matrix was used for downstream differential expression analysis, which were conducted as described above.

### DLX3 shRNA Transduction

After identifying that *DLX3* may be a cell state regulating transcription factor which is a target of DDE, we next wanted to assess the impact of DDE exposure in the context of *DLX3* knockdown. A *DLX3* shRNA lentiviral vector (Vector ID: VB250612-1662tnb) and a scrambled shRNA vector (Vector ID: VB010000-9548fjf) containing antibiotic resistance to hygromycin were built by VectorBuilder and lentiviral particles were synthesized. For lentiviral transduction, MCF10A cells were first cultured at 100,000 cells per well in a 6-well plate (Corning, #353046). After 24 hours, 5 µL of the 10^8^ TU/mL viral solution in 1 mL of cell culture media was added to two wells per condition. Following 48 hours of viral transduction, the virus-containing media was removed and 1 mL of media containing 25 μg/mL hygromycin (Gibco, #10687010) was added to each well. The cells were incubated in hygromycin-containing media for five days and split when needed once the wells reached 80-90% confluence. EGFP expression of the vector control was monitored to assess transduction efficacy. After five days of antibiotic selection, fresh media without antibiotics was added and the cells were grown until confluence. Cells were collected once EGFP expression was confirmed in the majority of the vector control-transduced cells via fluorescence microscopy.

To assess if we had generated a stable *DLX3* knockdown, first, RNA was extracted from non-transduced MCF10A cells, cells transduced with *DLX3* shRNA, and cells transduced with the scrambled shRNA. cDNA conversions were completed on extracted RNA using the iScript cDNA Synthesis Kit (BioRad, #1708890) according to the manufacturer’s protocol. Converted cDNA was combined with SYBR green master mix (BioRad, #1725270) and *DLX3*, *β-ACTIN*, or *GAPDH* primers (BioRad, qHsaCED0002786, qHsaCED0036269, qHsaCED0038674) and qPCR was run on a CFX96 Touch Real-Time PCR Detection System following the manufacturer’s protocol with three replicates per condition. *DLX3* knockdown was assessed by importing Cq values into R and calculating relative *DLX3* expression percentages between each condition, normalized by reference gene expression.

### DLX3 Knockdown Cell Culture, Treatment, and RNA-seq

Non-transduced MCF10A cells, cells transduced with the *DLX3* shRNA as well as the scrambled shRNA were cultured. Once all three cell types reached 80-90% confluency, the cells were collected and plated in mammosphere forming conditions with 25 µM DDE or 0.5% DMSO for seven days. Mammospheres from three replicates of each condition were collected, lysed, and stored in -80 °C until RNA extractions could be performed. *DLX3* expression was assessed via qPCR across non-transduced, *DLX3* shRNA transduced, and scrambled transduced mammospheres exposed to 25 µM DDE or control as described above. To perform a comprehensive assessment of the effects of DDE and/or *DLX3* knockdown, we quantified RNA using the Quant-it RiboGreen Reagent and RNA Assay Kit (Thermo, R11490) using 240 ng of RNA input and then performed RNA-seq using the MERCURIUS BRB-seq (Alithea Genomics, #10801) protocol as described in the Supplemental Methods with an input of 240 ng of purified RNA.

### Differential Gene Expression Analysis and Pathway Enrichment Analysis of DDE and DLX3 Knockdown

Differential gene expression was conducted the same as above with the following four comparisons: mammospheres transduced with the scrambled shRNA and exposed to 25 µM DDE (Scrambled-DDE) versus the scramble unexposed spheres (Scrambled-Control), mammospheres transduced with the *DLX3* shRNA and exposed to 25 µM DDE (DLX3-KD-DDE) versus the DLX3-KD unexposed spheres (DLX3-KD-Control), DLX3-KD-DDE versus Scrambled-DDE, and DLX3-KD-Control versus Scrambled-Control. We also analyzed the type-by-chemical interaction term (type: DLX3-KD vs Scrambled, chemical: DDE vs control) to determine if DDE impacted differential gene expression differently in the DLX3-KD spheres versus the Scrambled spheres. Pathway analysis was performed with the fgsea package v1.32.4 (29) and REACTOME database in R. Genes included in the analyses across the four comparisons and in the type-by-chemical interaction were ranked in descending order of logFC and analyzed for normalized enrichment scores in REACTOME pathways.

### Assessing DLX3 Expression in Breast Cancers from the Cancer Genome Atlas

To understand expression patterns of *DLX3* in normal breast and breast cancer, we next examined data from the Cancer Genome Atlas. UALCAN (30,31), a web browser that provides easy access to The Cancer Genome Atlas (TCGA) data, was used to assess *DLX3* transcripts per million in the breast invasive carcinoma dataset. *DLX3* expression was compared between normal breast tissue samples and samples from patients with breast invasive carcinoma with luminal, HER2-positive, and triple-negative cancer subtypes. Gene expression was also compared between normal TCGA samples and samples from cancer patients who self-report as European-American, African-American, and Asian-American. An unpaired t-test was used to evaluate levels of significance in *DLX3* expression level between any two sample groups.

## Results

### DDE Exposure Affects Stem Cell Proliferation

The cells capable of mammosphere formation are enriched for stem cell-like behavior and can undergo limited self-renewal (32). The mammosphere assay is therefore an important tool that provides readouts of breast stem cell anoikis resistance and proliferation capacity based on mammosphere count and size, as each mammosphere contains on average one sphere-forming stem cell. There were no significant changes in average mammosphere count with DDE exposure (Figure S1). Mammospheres exposed to 0.25 µM DDE had a significantly increased average area relative to control (p = 0.014), indicating an increase in cell proliferation (Figure S1).

### DDE Exposure Perturbs Mammary Epithelial Cell Identity and Stemness Markers

Next, we assessed the effects of DDE exposure on markers of cell identity and stemness, which we quantified via immunofluorescent high content imaging of mammospheres following KRT8 and KRT14 or CD44, CD24, and ALDH1A3 immunostaining. Representative immunofluorescent images are shown in Figure S2. The mean expression of KRT8 significantly decreased in mammospheres exposed to 25 µM DDE relative to control (p = 0.005)(Figure 1A). The mean expression of KRT14 was also significantly lower in DDE spheres exposed to 25 µM compared to control (p = 0.004). We did not observe significant changes in the intensity of CD24 or ALDH1A3 at any concentration of DDE (Figure 1B). As the concentration of DDE increased, the estimated mean expression of CD44 increased, with spheres exposed to 2.5 and 25 µM DDE having significantly higher intensity relative to control (p = 0.014 & 0.009, respectively) (Figure 1B).

**Figure 1.**
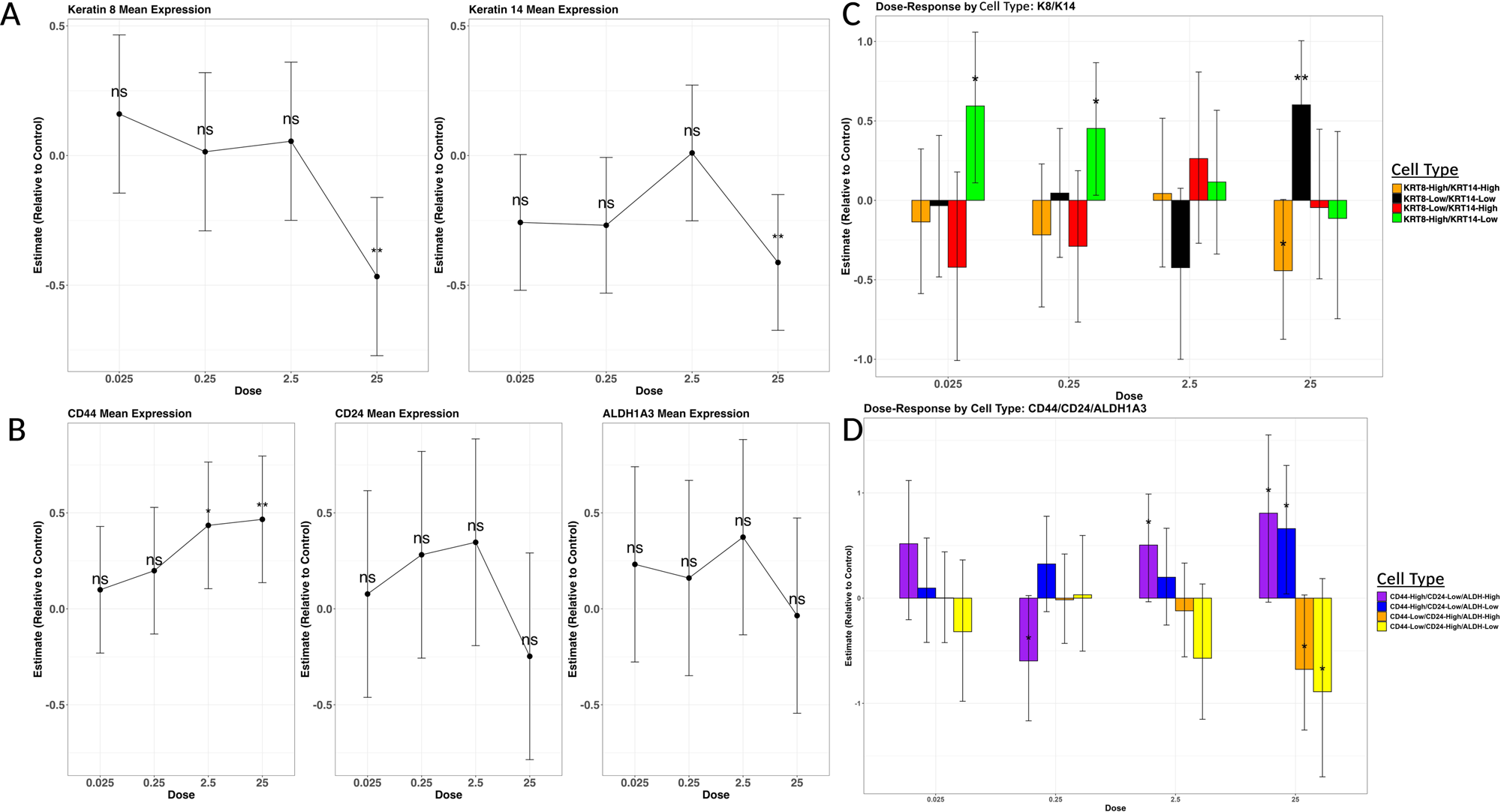
Generalized linear models of average immunofluorescent marker expression and proportions of cell phenotypes. A) Generalized linear models represent the robust normalized (median/median absolute deviation) and transformed (generalized log) estimated intensity units averaged per well for KRT8 and KRT14 across four concentrations of DDE compared to control and across three experimental rounds. Points represent model-estimated mean intensity values, and error bars represent approximate 95% confidence intervals (estimate ± 1.96 × standard error). B) Generalized linear models represent the robust normalized (median/median absolute deviation) and transformed (generalized log) estimated intensity units averaged per well for CD44, CD24 and ALDH1A3 across four concentrations of DDE compared to control and across three experimental rounds. Points represent model-estimated mean intensity values, and error bars represent approximate 95% confidence intervals (estimate ± 1.96 × standard error). C) Cell-type proportions were calculated after robust normalization (median/median absolute deviation) to the control group where cells were then classified as high or low for KRT8 and KRT14 relative to the control median. Bars represent the posterior mean estimate (c_effect) of the cell-type composition parameter for DDE-exposed samples relative to controls, based on three experimental rounds. Error bars indicate the 95% posterior credible interval, defined by the 2.5% (c_lower) and 97.5% (c_upper) quantiles of the posterior distribution. D) Cell-type proportions were calculated after robust normalization (median/median absolute deviation) to the control group where cells were then classified as high or low for CD44, CD24 and ALDH1A3 relative to the control median. Only a subset of populations were included for reporting (full Figure S4). Bars represent the posterior mean estimate (c_effect) of the cell-type composition parameter for DDE-exposed samples relative to controls, based on three experimental rounds. Error bars indicate the 95% posterior credible interval, defined by the 2.5% (c_lower) and 97.5% (c_upper) quantiles of the posterior distribution. P-values < 0.05 = *, < 0.01 = **, and < 0.001 = ***. P-values greater than 0.05 = ns.

After observing changes in expression of individual cell state markers by DDE exposure, we next wanted to evaluate whether DDE induced changes in the proportions of cells which jointly express cell state and stemness markers. To accomplish this, we classified single cells based on their cell state marker gene expression patterns and compared the proportion of cells in each cell state by exposure group. Figure S3 characterizes each cell population highlighted in the results. 0.025 µM and 0.25 µM DDE exposure shifted the cell population to have a greater estimated proportion of KRT8-High/KRT14-Low cells than in unexposed mammospheres (p = 0.024 & 0.046, respectively) (Figure 1C). Because the proportions are compositional, there was a non-significant but corresponding decrease in the proportion of KRT8-High/KRT14-High and basal-like KRT8-Low/KRT14-High cells in mammospheres exposed to 0.025 µM and 0.25 µM DDE compared to control. Surprisingly though, in mammospheres exposed to 25 µM DDE, the estimated proportion of KRT8-High/KRT14-High cells was significantly lower and the estimated proportion of KRT8-Low/KRT14-Low cells was significantly higher compared to control (p = 0.035 & 0.010, respectively), indicating reduced expression of both epithelial cell identity markers.

Next, we assessed differences in cell-type proportions based on single-cell stemness marker expression in mammospheres stained concurrently for CD44, CD24, and ALDH1A3 by DDE exposure (Figure 1D). Figure S3 describes each cell population highlighted in the results. All other CD44/CD24/ALDH1A3 cell population results are displayed in Figure S4. In mammospheres exposed to 0.25 µM DDE, the estimated proportion of CD44-High/CD24-Low/ALDH1A3-High cells (markers which reflect a hybrid epithelial-mesenchymal like stem cell) was significantly lower compared to control (p = 0.037), while in mammospheres exposed to 2.5 µM and 25 µM DDE, the proportion of these cells increased (p = 0.038 and p = 0.041, respectively). Additionally, in mammospheres exposed to 25 µM DDE, the estimated proportion of CD44-High/CD24-Low/ALDH1A3-Low cells was significantly higher, the estimated proportion of CD44-Low/CD24-High/ALDH1A3-High cells was significantly lower, and the estimated proportion of CD44-Low/CD24-High/ALDH1A3-Low cells was significantly lower compared to control (p = 0.036, 0.044, & 0.049, respectively).

In summary, 0.25 μM DDE exposure shifts the cell-type composition to have lower proportions of CD44-High/CD24-Low/ALDH1A3-High cells compared to control. However, 2.5 and 25 μM DDE have the opposite response, with greater proportions of the hybrid epithelial-mesenchymal stem cell-like CD44-High/CD24-Low/ALDH1A3-High populations. Regardless of ALDH1A3 expression, 25 μM DDE exposure also has a greater CD44-High/CD24-Low/ALDH1A3-Low population and corresponding decreases in CD44-Low/CD24-High/ALDH1A3-High epithelial stem-like cell populations and CD44-Low/CD24-High/ALDH1A3-Low more differentiated epithelial cell populations, compared to control. At 25 μM DDE, we also observed a loss of both KRT8 and KRT14 expression, suggesting that these cells were losing both their luminal and basal epithelial character.

### Transcriptomic Analyses Identify That DDE Induces a Signature of Epidermal Differentiation

To follow up on the findings on DDEs effects on markers of cell state and stemness, we next performed RNA-seq on mammospheres exposed to 0.025, 0.25, 2.5, 25 µM DDE or control for seven days to assess for differential gene expression and pathway enrichment attributable to DDE exposure. We found 585 genes that were significantly upregulated and 631 genes were downregulated in 25 µM DDE-exposed spheres compared to control out of 14,077 total genes (Figure 2A). Gene expression alterations with 25 µM DDE exposure were large: 33 of those genes significantly upregulated or downregulated had an absolute fold-change greater than 16, with four of those genes having absolute fold changes greater than 256 (*SMIM, H4C12, CUX2, LORICRIN*) (Figure 2B). In 2.5 µM DDE-exposed spheres, 21 genes were upregulated and 81 genes were downregulated compared to control. There were no significant differences in gene expression in 0.25 and 0.025 µM DDE-exposed spheres relative to control. A supplementary Excel file is available for each differential gene expression comparison (Data S1).

**Figure 2.**
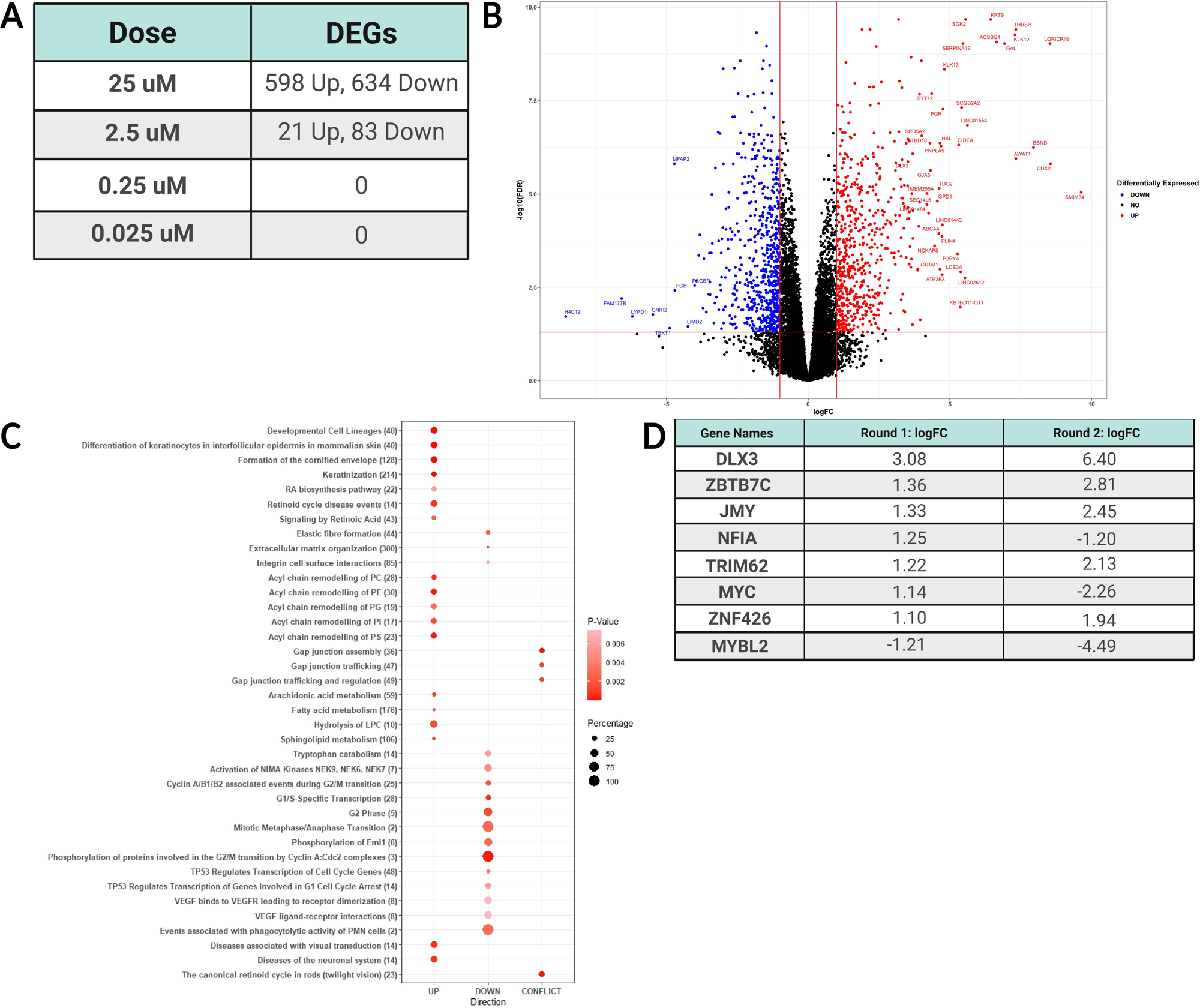
RNA sequencing analysis of mammospheres exposed to DDE. A) Table of the number of significantly differentially expressed genes (FDR adjusted p-value < 0.05 and |logFC| > 1) across four concentrations of DDE. B) Volcano plot of differentially expressed genes in 25 µM DDE-exposed mammospheres versus control. Points in red are upregulated genes compared to control and points in blue are downregulated genes compared to control. Points in black are genes that had a |log fold-change| less than 1 and/or an FDR adjusted p-value greater than 0.05. *DLX3* and all other genes with |logFCs| greater than 4 are labeled with their respective symbols. D) Dot plot of REACTOME pathway analysis comparing mammospheres exposed to DDE to unexposed mammospheres. REACTOME pathway names are listed on the y-axis with the total number of genes in that pathway listed in parentheses. The x-axis divides the pathways that are upregulated, downregulated, or have conflicting directions of differentially expressed genes. The percentage of genes differentially expressed out of the total number of genes in the pathway is depicted by the size of the dot and only pathways with a percentage equal to or higher than 10% and an FDR adjusted p-value less than 0.01 were included. FDR adjusted p-value is represented by dot color, with values closer to 0.05 shown as light red and values further from 0.05 as dark red. D) Table of genes identified as transcription factors and their respective logFCs that were significantly differentially expressed (FDR adjusted p < 0.05 and |logFC| > 1) in 25 µM DDE-exposed mammospheres compared to control in two rounds of RNA sequencing.

We then used BMDExpress 3 to quantify DDE exposure-response relationships of differential gene expression. Of interest, *LORICRIN*, a major component of the cornified cell envelope found primarily in the cells of the epidermis (33), had a model average benchmark concentration (BMC) of 10.80 µM and a fold-change around 512 at 25 µM. *DLX3,* a transcription factor that regulates epidermal cellular differentiation (34), had a model average BMC of 4.29 µM and a fold-change around 32 at 25 µM. We used the REACTOME database to conduct pathway enrichment analysis. 133 REACTOME pathways out of 1,347 were significantly differentially expressed in DDE-exposed spheres compared to control (Figure 2C). Pathways of interest that were significantly upregulated included “Formation of the cornified envelope”, “Keratinization”, and “Differentiation of keratinocytes in interfollicular epidermis in mammalian skin” (p = 3.09e-21, 2.44e-15, & 1.06e-07, respectively). These terms had BMC means between 7.19 and 9.13 µM and mean fold changes between 5.64 and 6.83. These RNA-seq results suggest that the higher concentrations of DDE are inducing a squamous epidermal-like cellular phenotype.

### Cardamonin Concentration-Dependently Attenuates the Effects of DDE on Epidermal Differentiation Signature

In a previous experiment, we had identified that the phenotypic effects of DDE mimicked that of the Wnt pathway activator CHIR99021 and that DDE exposure in MCF10A cells induces translocation of the canonical Wnt signaling pathway member β-catenin (26). Given established links between Wnt activation and squamous epidermal metaplasia in the breast (35), we next wanted to assess whether DDE’s effects on the induction of epidermal cell markers was linked to Wnt signaling. We tested the effects of cardamonin, a known Wnt inhibitor (27), alone and in combination with DDE to determine if DDE was acting through a Wnt activation pathway to induce epidermal differentiation of the mammospheres. We found that cardamonin had no detectable cytotoxicity at 3.75 µM and 0.94 µM but there were significant decreases in β-catenin expression after 48 hours of 3.75 µM cardamonin exposure, consistent with Wnt pathway inhibition (Figure S5). Therefore, we chose these two concentrations to test interactions between DDE and cardamonin. We identified 526 significantly differentially expressed genes (compared to control) when comparing spheres exposed to 25 µM DDE alone and the mixtures of 25 µM DDE and cardamonin (Figure S6A). We then used BMDExpress 3 to investigate REACTOME pathway enrichment. By comparing 25 µM DDE, 25 µM DDE + 0.94 µM cardamonin, and 25 µM DDE + 3.75 µM cardamonin to assess the effects of increasing cardamonin, we found downregulation of the pathways “Differentiation of keratinocytes in interfollicular epidermis in mammalian skin” and “Formation of the cornified envelope” (p = 0.002 & 0.001, respectively; Figure S6B), indicating that in a mixture of DDE and cardamonin, increasing concentration of cardamonin can attenuate effects of DDE on epidermal differentiation. Next, we compiled the list of genes that were differentially expressed in the REACTOME pathways “Keratinization”, “Formation of the cornified envelope”, and “Differentiation of keratinocytes in interfollicular epidermis in mammalian skin” in at least one condition compared to control. When comparing skin-related gene expression between cardamonin and mixtures of 25 µM DDE and cardamonin or 25 µM DDE alone, cardamonin exposure flipped the fold-change direction of the genes *PERP* and *DSG1* (Figure S6C). *PERP* is a p53 target gene involved in DNA damage-induced apoptosis (36). *DSG1* encodes a member of the desmoglein protein subfamily which are major components of the desmosome (37). When comparing mixtures of 25 µM DDE and cardamonin with 25 µM DDE alone, the addition of cardamonin did not flip the fold-change direction of any gene but did reduce the fold-change magnitude and significance of genes such as *LORICRIN*, *DLX3*, late cornified envelope genes, and filaggrin genes. However, only two skin-related genes were significantly upregulated with 2.5 µM DDE exposure. When comparing mixtures of 2.5 µM DDE and cardamonin with 2.5 µM DDE alone, the addition of cardamonin actually induced differential gene expression of many skin-related genes, including *DLX3* and many late cornified envelope genes. We also tested CHIR99021, a Wnt activator, as a Wnt signaling positive control. CHIR99021 exposure induced differential gene expression of some skin-related genes including the upregulation of *DLX3*, but at a much lower magnitude than 25 µM DDE exposure, compared to control (Figure S6C).

Overall, cardamonin exposure alone can downregulate some skin-related genes that are upregulated in the presence of DDE. However, when in a mixture with 25 µM DDE, cardamonin only modestly reduces the magnitude of differential expression among epidermal differentiation markers. Conversely, a mixture of 2.5 µM DDE and cardamonin can induce expression of skin-related genes that 2.5 µM DDE alone does not, indicating that cardamonin may have concentration-dependent effects of attenuation or potentiation of epidermal differentiation. Additionally, CHIR99021 treatment recapitulated the induction of some skin-related genes but at a much lower magnitude, suggesting that the effects of DDE on the metaplastic phenotype may include Wnt signaling but other signaling pathways are involved (Data S1).

### DDE Induces Epidermal Differentiation through DLX3-dependent and Independent Signaling

After only modest impacts on the epidermal differentiation signature was observed after cardamonin treatment, we reexamined the DDE exposure gene expression signatures from both rounds of RNA-seq to identify if there may be other potential drivers of the DDE induced epidermal gene expression signature. We identified eight transcription factors in both rounds of RNA sequencing which had significant differential gene expression in mammospheres exposed to 25 µM DDE compared to control (Figure 2D). Of those, we prioritized *DLX3* for additional experimental investigation due to having the highest average fold-change between the two rounds as well as its role in the identified epidermal differentiation-related REACTOME pathways that were significantly upregulated in DDE-exposed spheres.

We constructed a lentiviral *DLX3* shRNA for stable knockdown of *DLX3* expression to determine if DDE-induced epidermal differentiation upregulation would be inhibited by *DLX3* knockdown. After successful knockdown in MCF10A cells (Figure S7A), we exposed MCF10A cells transduced with the *DLX3* shRNA or a scrambled shRNA to 25 µM DDE or control for seven days in mammosphere-forming conditions. We collected these mammospheres and confirmed sustained *DLX3* knockdown (Figure S7B), and performed RNA-seq to understand how DDE exposure impacts mammospheres in control and *DLX3* knockdown conditions.

We conducted differential gene expression analysis first comparing DDE and control exposed scramble shRNA-treated mammospheres. We found that 231 genes were significantly upregulated and 174 genes were significantly downregulated out of 10,544 total genes with DDE exposure (Figure S8A). The differential gene expression and REACTOME pathway enrichment (Figure S8B) patterns reflected what we observed in RNA-seq of the non-transduced mammospheres, including strong induction of epidermal markers such as *LORICRIN*. This indicated that the scramble shRNA-treated mammospheres had a comparable phenotype to non-transduced mammospheres with respect to effects of DDE exposure.

Differential gene expression analysis comparing DLX3-KD-DDE mammospheres and Scrambled-DDE mammospheres showed 76 significantly upregulated genes and 90 significantly downregulated genes (Figure 3A). As hypothesized, *DLX3*, *LORICRIN*, and REACTOME pathways such as “Keratinization” and “Formation of the cornified envelope” were significantly downregulated in DLX3-KD-DDE mammospheres compared to Scrambled-DDE mammospheres (Figure 3A & 3B). This suggests that DLX3 is important for regulating the induction of the squamous metaplasia keratinization gene expression signature following DDE exposure.

**Figure 3.**
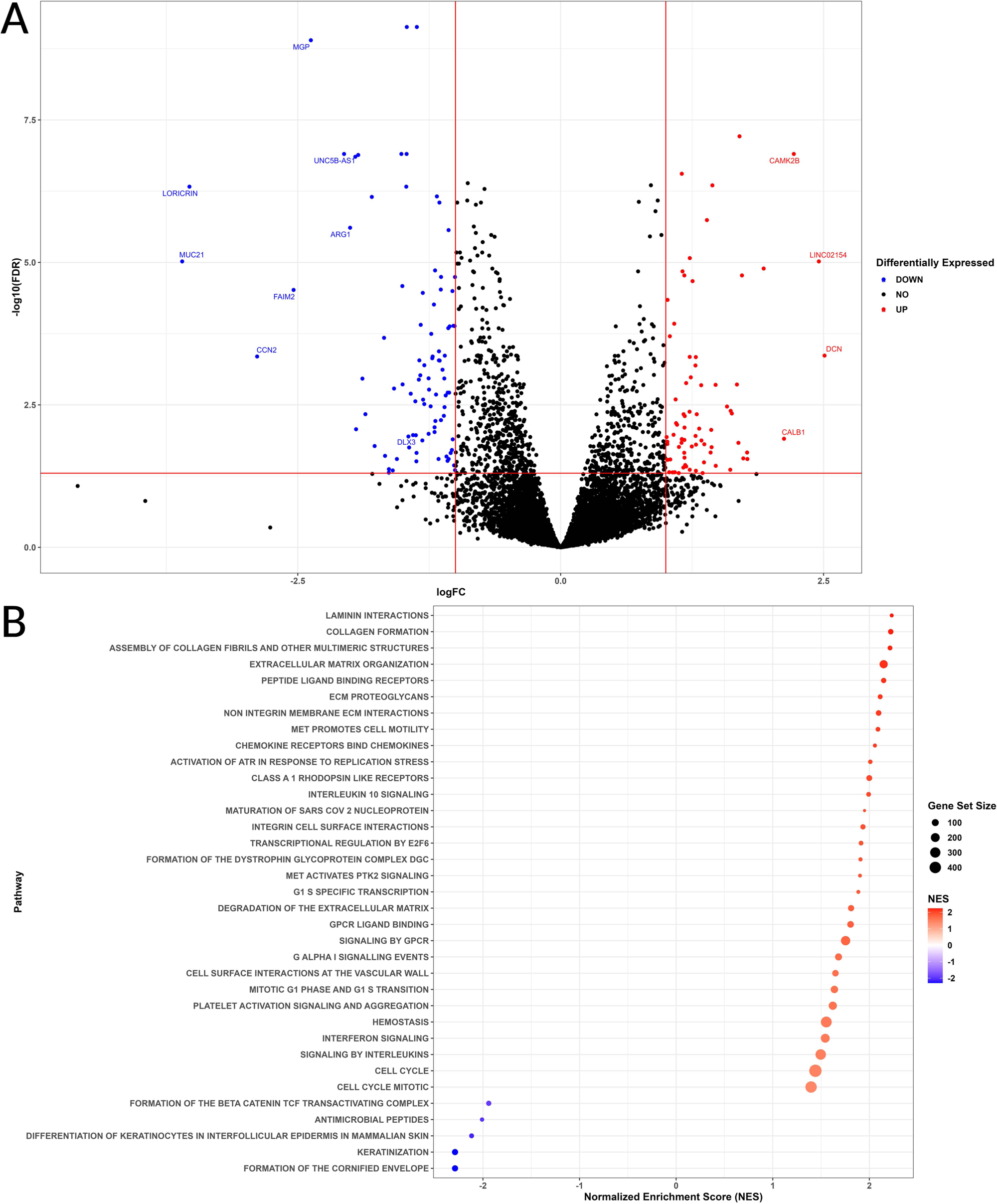
RNA sequence analysis of DLX3-KD-DDE (25 µM) mammospheres versus Scrambled-DDE (25 µM) mammospheres. A) Volcano plot of differentially expressed genes. Points in red are upregulated genes compared to control and points in blue are downregulated genes compared to control. Points in black are genes that had a |log fold-change| less than 1 and/or an FDR adjusted p-value greater than 0.05. *DLX3* and all other genes with |logFCs| greater than 2 are labeled with their respective symbols. B) Dot plot of REACTOME pathway analysis comparing DLX3-KD-DDE mammospheres versus Scrambled-DDE mammospheres. REACTOME pathway names are listed on the y-axis. The x-axis separates upregulated pathways with a positive normalized enrichment score (red) and downregulated pathways with a negative normalized enrichment score (blue). The darker shaded points represent higher absolute scores. The size of the dot represents the total number of genes in the pathway. Only pathways with an FDR adjusted p-value less than 0.05 were plotted.

Next, we compared DLX3-KD-DDE mammospheres versus DLX3-KD-Control mammospheres, to further assess how DDE exposure impacts gene expression in the presence or absence of DLX3 knockdown. We identified 143 upregulated genes and 125 downregulated genes between the conditions (Figure S9A). Genes such as *LORICRIN* and related REACTOME pathways such as “Keratinization” and “Formation of the cornified envelope” (Figure S9B) were upregulated in DLX3-KD-DDE mammospheres vs DLX3-KD-Control. DLX3-KD-Control mammospheres had a 21.8% decrease in *DLX3* counts per million (CPMs) and a 97.7% decrease in *LORICRIN* CPMs (Figure S9C) compared to Scrambled-Control. DDE exposure in mammospheres transduced with the *DLX3* shRNA restored *DLX3* and *LORICRIN* gene expression levels comparable to those found in Scrambled-Control. In comparison, DDE exposure in mammospheres transduced with the scrambled shRNA had a 204.6% increase in *DLX3* expression and a 1609.8% increase in *LORICRIN* expression. This suggests that DDE can still induce *DLX3* expression in the presence of DLX3 shRNA, but at a decreased amount compared to control cells.

Differential gene expression analysis assessing the interaction between DLX3-KD and DDE exposure identified 19 significantly upregulated genes and 78 significantly downregulated genes which were significantly associated with the interaction (Figure S10A). REACTOME pathways relating to glucose metabolism were significantly downregulated and a pathway related to transcription was significantly upregulated in the interaction (Figure S10B). While statistical significance was not reached, the effect of DDE was more negative for *DLX3* expression in DLX3-KD spheres compared to Scrambled spheres. Other skin-related genes were significantly differentially impacted by DDE between DLX3-KD spheres compared to Scrambled spheres (Figure S10C).

Overall, *DLX3* knockdown was effective in downregulating both *DLX3* and expression of keratinization associated genes, like *LORICRIN*. DDE exposure did not have a significant differential effect between DLX3-KD spheres and Scrambled spheres for these genes. However, other skin-related genes were differentially impacted, which could indicate that the *DLX3* knockdown was not complete or that DDE acts through both *DLX3*-dependent and *DLX3*-independent mechanisms to induce epidermal differentiation.

### Changes in DLX3 Expression Yields Distinct Pathway Enrichments and Outcomes in Breast Cancer Patients

As our previous experiments suggested that *DLX3* is a transcription factor which is inducible by DDE and regulates cell state transitions, we next wanted to understand the effects of *DLX3* expression itself in breast epithelial cells. We compared gene expression in the DLX3-KD mammospheres versus Scrambled shRNA mammospheres, both unexposed to DDE, where we identified 154 upregulated and 113 downregulated genes (Figure 4A). Noteworthy REACTOME pathways upregulated following DLX3-KD involve interferon signaling, interleukin signaling, and extracellular matrix (ECM) interactions and organization (Figure 4B). We next wanted to understand DLX3 expression in breast cancer compared to normal breast tissue. By using UALCAN to interrogate the Cancer Genome Atlas (TCGA), we compared *DLX3* transcripts per million (TPMs) between normal breast tissues and luminal, HER2 positive, and triple negative breast cancers (Figure 4C). *DLX3* was increased in all three cancer subtypes compared to normal patients (p = <1.00e-12, 2.60e-04, & 1.10e-06, respectively). There were no significant differences in *DLX3* TPMs when comparing within the three cancer subtypes. We also compared *DLX3* TPMs between normal patients and breast cancer patients who are European-American, African-American, and Asian-American (Figure 4D). *DLX3* was also upregulated in tumors from European American, African American, and Asian American breast cancers compared to normal tissues (p = <1.00e-12, 3.67e-08, & 9.96e-05, respectively). Of note, *DLX3* was significantly higher in African-American cancer patients compared to European-American cancer patients (p = 0.044).

**Figure 4.**
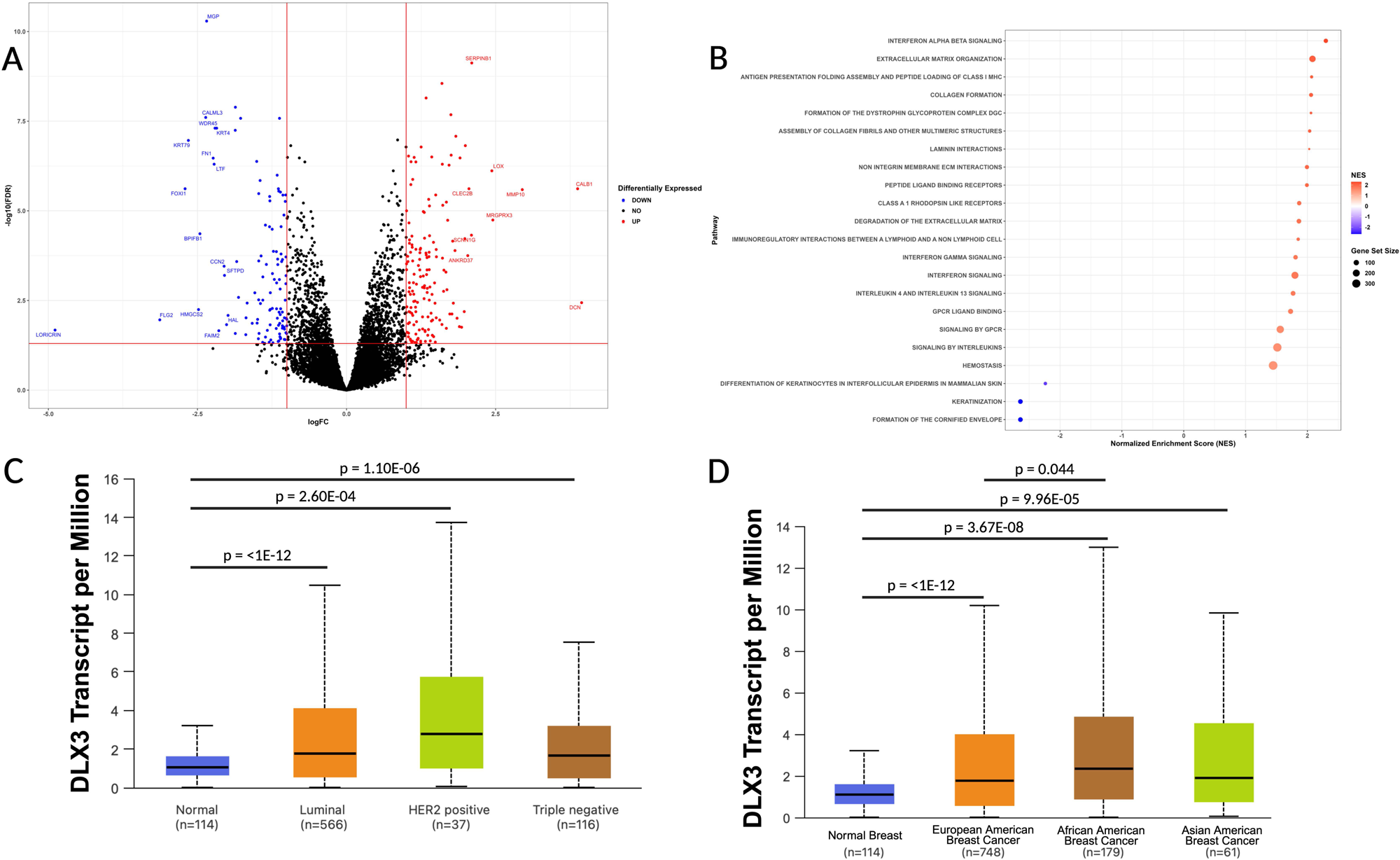
RNA sequence analysis of DLX3-KD-Control mammospheres versus Scrambled-Control mammospheres and TCGA analysis of *DLX3* transcripts per million. A) Volcano plot of differentially expressed genes. Points in red are upregulated genes compared to control and points in blue are downregulated genes compared to control. Points in black are genes that had a |log fold-change| less than 1 and/or an FDR adjusted p-value greater than 0.05. Genes with |logFCs| greater than 2 are labeled with their respective symbols. B) Dot plot of REACTOME pathway analysis comparing DLX3-KD-Control mammospheres versus Scrambled-Control mammospheres. REACTOME pathway names are listed on the y-axis. The x-axis separates upregulated pathways with a positive normalized enrichment score (red) and downregulated pathways with a negative normalized enrichment score (blue). The darker shaded points represent higher absolute scores. The size of the dot represents the total number of genes in the pathway. Only pathways with an FDR adjusted p-value less than 0.05 were plotted. C & D) Boxplots with *DLX3* transcripts per million compared across normal patients (blue) and patients with breast invasive carcinoma. The number of patients in each group is listed on the x-axis. An unpaired t-test was used to evaluate levels of significance in *DLX3* expression level between any two sample groups. P-values with significant comparisons are shown above the boxplots with bars spanning the two groups being compared. C) Three cancer subtypes, luminal, HER2 positive, and Triple negative, are represented by orange, green, and brown boxplots, respectively. D) Three cancer patient races, European-American, African-American, and Asian-American, are represented by orange, brown, and green, respectively.

## Discussion

Breast cancer is one of the most common cancers worldwide (38). Increasing evidence shows that alongside genetic and socioeconomic factors, environmental factors play a large role in breast cancer risk (39). Both epidemiological and mechanistic studies show associations between DDT exposure and breast cancer (5,40). DDT and its metabolite DDE have long half-lives via bioaccumulation in body fat, indicating that humans may still be at risk of harm even after prohibition of DDT in the 1970s (40). With inconsistencies about the associations between DDE and breast cancer risk, we aimed to assess the effects of DDE exposure on breast cell morphology, phenotypic plasticity, and gene expression. This research identified perturbations in cell identity, stemness, and gene expression related to epidermal differentiation in non-tumorigenic breast epithelial cells grown as mammospheres in three dimensional non-adherent tissue culture conditions, identifying a novel mechanism by which environmental exposures may promote cellular plasticity and breast carcinogenesis.

Immunofluorescent analysis of cell state marker protein expression showed distinct changes at each concentration level of DDE. Following exposure to 0.025 and 0.25 µM DDE, the cell-type composition shifted to more KRT8-High/KRT14-Low cells, indicating a more luminal state. At 2.5 µM DDE, there was a greater proportion of CD44-High/CD24-Low/ALDH-High cells and an increase in CD44 intensity compared to control, signifying that these cells are expressing proteins associated with a hybrid epithelial/mesenchymal phenotype. Lastly, 25 µM DDE shows a decrease in KRT8 and KRT14 expression and an increase in CD44-High/CD24-Low cell populations, demonstrating that these cells are losing their mammary luminal and basal epithelial cell identity markers and expressing more CD44, consistent with a more mesenchymal stem-like state. These findings indicate a non-monotonic response, where nanomolar concentrations of DDE shifted cell populations to a more luminal phenotype, but 25 µM DDE exposure conferred a more mesenchymal stem-like marker expression phenotype.

After conducting RNA-seq analysis, the most striking genes and enriched pathways found to be differentially expressed following DDE exposure were those related to epidermal differentiation, keratinization, and cornification. These enriched pathways are features of keratinizing squamous cell metaplasia (41). Breast squamous metaplasia is a rare but observed phenotype in typically benign fibroadenomas, cysts, abscesses, and chronic inflammation (42). *In vivo* experiments have found cornification and epidermal differentiation genes upregulated in mouse models mimicking human triple-negative breast cancer via inactivation of *Rb*, *p53*, and *Brca1* in the mammary gland (43). Toxic and mechanical injury have been reported to induce squamous metaplasia in a number of other tissues (41). For example, cigarette byproducts can cause localized hypoxia or tissue damage to induce keratinizing squamous metaplasia in the breast (44). Here, we report that exposing MCF10A-derived mammospheres to 25 µM DDE is a reliable method to induce a squamous cell metaplasia gene expression phenotype *in vitro*.

To understand the mechanism behind the epidermal differentiation signature, we tested if this process was occurring via Wnt signaling and could be counteracted with co-exposure to DDE and the Wnt inhibitor cardamonin (37). When treated with a combination of 25 µM DDE and cardamonin, increasing concentration of cardamonin can attenuate effects of DDE on epidermal differentiation, however, this attenuation is modest. Conversely, when treated with a combination of 2.5 µM DDE and cardamonin, the cardamonin actually induces the effects on epidermal differentiation compared to 2.5 µM DDE alone, indicating a non-monotonic effect of the mixtures. Moreover, treatment with CHIR99021, a Wnt activator, recapitulated the increased expression of some epidermal genes but at a much lower magnitude than DDE. The effects of DDE on the metaplastic phenotype may include Wnt signaling but other signaling pathways are likely also involved.

Through additional mining of our RNA-seq data, we identified *DLX3* as a potential transcription factor associated with epidermal differentiation of DDE-exposed mammospheres. *DLX3* is a transcription factor which drives skin development and generally thought of as a tumor suppressor gene that co-regulates the cell cycle with p53, where loss of *DLX3* function is associated with skin tumor progression (45). This mechanism is primarily studied in squamous cell carcinomas with a key research gap about *DLX3* expression in breast cancer, particularly given its ability to be induced by DDE exposure in this study. We tested if knockdown of *DLX3* would attenuate DDE’s effects on induction of a squamous metaplasia expression signature. As expected, *DLX3* and the pathways related to squamous cell metaplasia were significantly downregulated in the DLX3-KD DDE-exposed mammospheres compared to DDE-exposed mammospheres with normal *DLX3* expression. While it did not reach statistical significance, DDE had a more negative impact on *DLX3* expression in DLX3-KD spheres compared to

Scrambled spheres. The nonsignificance may stem from the incompleteness of the *DLX3* knockdown. In DLX3-KD spheres, baseline expression of *DLX3* was lower than Scrambled spheres, but we still observed significant upregulation of *DLX3* from DDE exposure in the context of the DLX3-KD. However, the level of DLX3 expression did not approach the maximal expression level reached by Scrambled-DDE spheres following DLX3 exposure. Furthermore, other skin-related genes were differentially impacted by DDE between the DLX3-KD and the Scrambled control. Thus, DDE likely acts through *DLX3*-dependent and *DLX3*-independent mechanisms in order to produce the epidermal differentiation signature.

When assessing the effect of *DLX3* knockdown in the absence of DDE exposure, pathways related to interferon signaling, interleukin signaling, and ECM interactions and organization were upregulated. Interferon and interleukin signaling pathways are important for tumor recognition and destruction through immunosurveillance (46,47). The ECM mediates communication between mammary epithelial and stromal cells, and is important for regulating cell proliferation, differentiation, and migration (48). In breast cancer, ECM degradation is required for cancer cell invasion and metastasis (49). If hypothesizing that overexpression of *DLX3,* such as in the context of DDE exposure, would downregulate these pathways, DDE exposure may create a more favorable environment for breast cancer development by suppressing immune function and promoting ECM remodeling.

In the TCGA database, *DLX3* expression was significantly elevated in tissues from breast cancer patients across all three cancer subtypes compared to patients without cancer. *DLX3* expression was also significantly higher in African-American versus European-American cancer patients. African-American women also experience higher incidence of triple-negative breast cancer and higher levels of DDE exposure compared to European-American women (9,50). Thus, inappropriate DLX3 overexpression may be a mechanism driving breast carcinogenesis which can be environmentally modulable by exposures such as DDE.

### Limitations and Future Directions

*In vivo,* estrogen-receptor (ER) α was found to be a regulator of Dlx3 function (51). While DDE is a known endocrine disruptor, the cell line used in our experiments, MCF10A, is ER-negative (52). This suggests that DDE may be inducing *DLX3* expression through an estrogen-independent mechanism in this cell model. Future work, however, could be conducted in ER-positive cells exposed to DDE to determine if there is an additional role of upstream estrogen-receptor signaling on *DLX3*-driven squamous cell metaplasia. Furthermore, we did not achieve 100% *DLX3* knockdown efficiency, leading to potential variability in our outcomes due to expression heterogeneity. Future work could conduct a full knockdown to mechanistically assess whether DDE’s induction of a squamous metaplasia expression signature requires *DLX3*. This research would also be moved forward by dissociating exposed mammospheres and implanting them into a mouse model to assess their tumorigenicity.

### Summary

In our study, we exposed mammospheres to human relevant concentrations of DDE and found significant perturbations of cell identity and stemness markers and further transdifferentiation of mammary epithelial cells to an epidermal cell phenotype. We also found that exposing MCF10A-derived mammospheres to 25 µM DDE is a reliable method to induce a squamous cell metaplasia expression profile *in vitro*. *DLX3* knockdown was effective in reducing the magnitude of DDE-induced epidermal differentiation. By investigating patient sample data, we found that *DLX3* expression is upregulated in breast cancer patients. Collectively, these findings could provide an additional mechanistic link between the increased risk of breast cancer in individuals disproportionately exposed to DDE.

## Acknowledgements

Funding for this study was provided by the University of Michigan Rogel Cancer Center Scholars Program and the National Institutes of Health (R01 ES028802, P30 ES017885, P30 CA046592).

