## Supplemental_Methods_and_Figures for "Exposure to p,p’-dichlorodiphenyldichloroethylene (p,p’-DDE) Perturbs Cell State and Stem Cell Markers and Induces an Epidermal Phenotype in Human Breast Cells"

*Cell Culturing and Mammosphere Formation*

For mammosphere culture experiments, five 6-well (Fisher, 07-200-601) and one 96-well ultra-low adherent plates (Fisher, 07-200-603) were used for plating mammospheres. 200,000 cells in 2 mL of DDE or control media were added to each well of the 6-well plates with five technical replicates for each treatment. 5,000 cells in 100 µL of DDE or control media were added to each well of the 96-well plate with six technical replicates of each treatment. The plates incubated at 37°C and 5% CO_2_ for seven days with 40 µL of DDE or control mammocult media to the 96-well plate and 1 mL to the 6-well plates added three days after plating.

*Quantification of Mammosphere Size and Number*

To quantify mammosphere number and size, spheres were stained with Hoechst 33342 (Fisher, H3570). The 96-well plate was knocked to ensure that mammospheres were centered in the well before adding 40 µL of the Hoechst 33342 solution (1:2000) to each well in the dark. The plate was wrapped in foil to protect from light and incubated at 37°C for 30 minutes. The stained plates were imaged using an Invitrogen EVOS M7000 imaging system software version 2.0.2094.0. Automated imaging was set up at 10x EVOS 8.4 objective. Exposure time was set based on control wells. All fields for each well were captured at a single z height with autofocus set at the stained channel with Hoechst (350/461). One raw stitched image per well was saved along with per-field raw images in .TIF format.

*Cell State and Stem Cell Marker Immunostaining High Content Imaging*

After 24 hours, the cells were fixed by carefully removing 95 µL of the media and adding 100 µL of 4% methanol-free paraformaldehyde (Thermo, AA433689M) prepared in PBS to each well. The plate was incubated for 20 minutes at room temperature. The plates were washed once with 1X PBS and permeabilized with 100 µL of 1% Triton-X 100 (Sigma Aldrich, T8787) prepared in 1X PBS for 20 minutes. Two blocking buffers were prepared for the KRT8/KRT14 (luminal and basal “keratin”) panel and the CD44/CD24/ALDH1A3 (epithelial and mesenchymal “stemness”) panel. For the keratin panel, plates were washed twice with 1X PBS and then incubated for 1 hour at room temperature with 100 µL per well of 1% bovine albumin fraction V (BSA) (Gibco, 50-121-5315) and 22.5 mg/mL glycine (Thermo, AAA1381636) in a solution of 1X PBS and 0.1% Tween 20 (PBST) (Fisher, BP337). For the stemness panel, the blocking buffer contained 1% BSA in PBS. The blocking solution was removed and 100 µL of the primary conjugated KRT8/KRT14 antibody solution (1:150/1:50)(Abcam, ab192467; Abcam, ab192056) prepared in 1% BSA in PBST was added to each well in the respective plate and 100 µL of the CD44 (primary conjugated)/CD24 (primary conjugated)/ALDH1A3 antibodies (1:625/1:25/1:312.5) (Biolegend, 103016; Santa Cruz, sc-19585 AF647; Abcam, ab129815) in 1% BSA in PBS was added to each well in the other plate. This was completed in the dark and the plates were wrapped in foil and incubated overnight at 4 °C.

After 8-12 hours, the staining solution was removed and the KRT8/KRT14 plate was washed three times with PBST and the CD44/CD24/ALDH1A3 plate with PBS in the dark to avoid stain bleaching. 100 µL of PBS/PBST containing Hoechst 33342 (DNA stain) was added to each plate and a goat anti-rabbit secondary antibody to ALDH1A3 (1:1000)(Abcam, ab150080) was added to the CD44/CD24/ALDH1A3 plate. After 1 hour incubation at room temperature, both plates were then washed with 100 µL of PBST three times. Three technical replicates per condition were set up for each staining panel. A Yokogawa CV8000 microscope was used for automated plate imaging with a ×20/1.0NA water immersion objective lens, 50-µm pinhole. Laser power for each channel was adjusted to ensure optimal signal-to-noise ratios and laser-based autofocus was performed at each field with 16 fields per well captured. Hoechst was imaged in the 405 nm channel, KRT8 and CD44 at 488 nm, ALDH1A3 at 594 nm, and KRT14 and CD24 at 647 nm.

*RNA-Sequencing Analysis of DDE Effects in Mammospheres*

Four replicates of mammospheres from each condition were collected and washed with 2 mL of normal mammocult media to ensure all spheres were collected off the plate. They were centrifuged at 110 g for 10 minutes and washed twice by resuspending spheres in 2 mL PBS. Spheres were then lysed with 350 μL of Buffer RLT (Qiagen, 80004) containing 1% β-mercaptoethanol (Sigma Aldrich, M6250). Lysates were stored at -80 C until extractions. Lysates were homogenized via QIAShredder columns (Qiagen, 79654) and RNA was extracted using the Qiagen AllPrep DNA/RNA/Protein Mini kit (Qiagen, 80004), following manufacturer’s protocols. RNA was quantified using the Quant-it RiboGreen Reagent and RNA Assay Kit (Thermo, R11490) using 10 ng/μL of RNA input. Extracted RNA was stored at -80C until library preparation and sequencing could be performed. cDNA and library preparation was performed using the SeqWell plexWell plate-based preparation kit, following our previously defined protocol (10). Libraries were sequenced on a Novaseq 6000 (Illumina) at the University of Michigan Advanced Genomics Core Facility. Sequencing read quality was assessed via FastQC (53) and MultiQC (54) prior to reads being aligned to a splice junction aware build of the human genome (GRCh38) using STAR (55). Aligned reads were assigned to genes via featureCounts (56), not counting multimapping and multi-overlapping reads.

*Cardamonin Cell Viability and Assaying β-catenin Expression*

MCF10A cells were seeded in a T-75 flask and allowed to grow at 37 °C and 5% CO2 until they reached 80% confluency. Cells were trypsinized and plated in two 384-well plates (Revity, 6057602) at 400 cells/well with six replicates per condition. The plates were incubated without treatment for 24 hours before being exposed to the chemical conditions to ensure that the cells had adhered. Cardamonin (Sigma Aldrich, 681672-2MG) was dissolved in DMSO and the following concentrations were prepared in MCF10A media: 30, 15, 7.5, 3.75, 1.88, 0.94, 0.47, 0.23, 0.12, and 0.06 µM. 0.5% DMSO in MCF10A media was used for the vehicle control. One plate was exposed to cardamonin on the first day and both plates were exposed 48 hours later. After one plate grew in cardamonin or control conditions for 96 hours and the other for 48 hours, both plates were fixed with 4% PFA in PBS for 20 minutes at room temperature. Then, the wells were washed once with PBS before adding 50 µL of 0.1% Triton-X 100 in PBS to permeabilize the cells. This solution sat for 15 minutes at room temperature. The plates were washed twice with PBS and then incubated for 1 hour at room temperature with 1% BSA and 0.1% Tween 20 in PBS. A 1:100 solution of β-catenin primary antibody (Thermo, 71-2700) in 1% BSA and 0.1% Tween 20 in PBS was added to each well and incubated overnight at 4 °C. The wells were washed three times with 0.1% Tween 20 in PBS. A 1:1000 solution of goat anti-rabbit secondary antibody to β-catenin (Abcam, ab150079) and the nuclear stain Hoechst 33342 in 1% BSA and 0.1% Tween 20 in PBS was added to each well for 1 hour. The wells were washed three times with 0.1% Tween 20 in PBS and the plates were imaged on a Thermo Scientific CellInsight CX5 microscope using HCS Studio: Cellomics Scan v6.6.1 at 20x objective with 9 fields per well. Hoechst imaging was captured in the 405 nm channel and β-catenin in the 647 nm channel.

*Drug-seq cDNA and Library Preparation*

Mammospheres were collected, washed with PBS, and flash frozen at -80 C before being prepared as lysates following the Alithea Genomics MERCURIUS Drug-seq protocol and kit (Alithea Genomics, #10701) using the 1x Cell Lysis Buffer with RNase Inhibitor (Alithea Genomics, #10351). Mammospheres were formed with 200,000 cells per sample but the Drug-seq protocol recommends a maximum input of 50,000 cells. To address this, spheres were lysed in 20 µL of the lysis buffer but only 5 µL of the lysate was used for library preparation. After the spheres were lysed, the samples were reverse-transcribed via mixing with the barcoded oligo-dT primers provided in the 96-well PCR plate. A reverse transcription master mix was then added to each well and a Bio-Rad PTC Tempo Deepwell Thermal Cycler was used to facilitate the reaction.

Next, i7 indexed samples were pooled into a 2 mL tube and cDNA was purified using an equivalent volume of SPRI beads (Beckman Coulter, A63881). The cDNA and added bead solution incubated for 5 minutes at room temperature and was then placed on a magnet (Thermo, 12321D) for 5 minutes. The supernatant was removed and the beads were washed twice with 80% ethanol (Decon Labs, 2716). The cDNA was eluted with 37 µL Dnase/RNase-free deionized water (Thermo, A57775). From there, 35 µL was transferred to a 0.2 mL PCR tube (Fisher, 14230225). Free primer digestion and second strand synthesis was then performed by using the manufacturer’s buffer and enzyme solutions and incubating on the Thermal Cycler. A 0.6x ratio of SPRI magnetic beads was used for double-stranded cDNA purification and beads were washed twice with 80% ethanol before eluting with 21 µL of water.

cDNA concentration was determined by a Picogreen cDNA assay (Thermo, P7589). A black clear-bottom 96-well plate was used and samples and standards were incubated with the Picogreen reagent before fluorescence was measured on a SpectraMax M5e plate reader. After the cDNA was quantified, a tagmentation mix was prepared with 20 ng of cDNA input. The tagmentation solution with added cDNA incubated for 7 minutes at 55C and was immediately put on ice. Tagmented cDNA was purified with a 0.6x ratio of SPRI beads and two washes of 80% ethanol. 20 µL of the eluted cDNA in water was transferred to a new tube.

Following tagmentation, the 5’ terminal fragments are amplified using the Unique Dual Indexing (UDI) adapter primers. A library amplification mix, the UDI adapter, and the tagmented cDNA are mixed and 14 cycles of PCR amplification were performed on the Thermal Cycler. Lastly, the cDNA library was purified with SPRI magnetic beads at a 0.7x ratio and 80% ethanol. Beads were washed twice and the cDNA was elution with water. The purification was performed twice to effectively remove primer dimer fragments.

**Supplemental Figures**

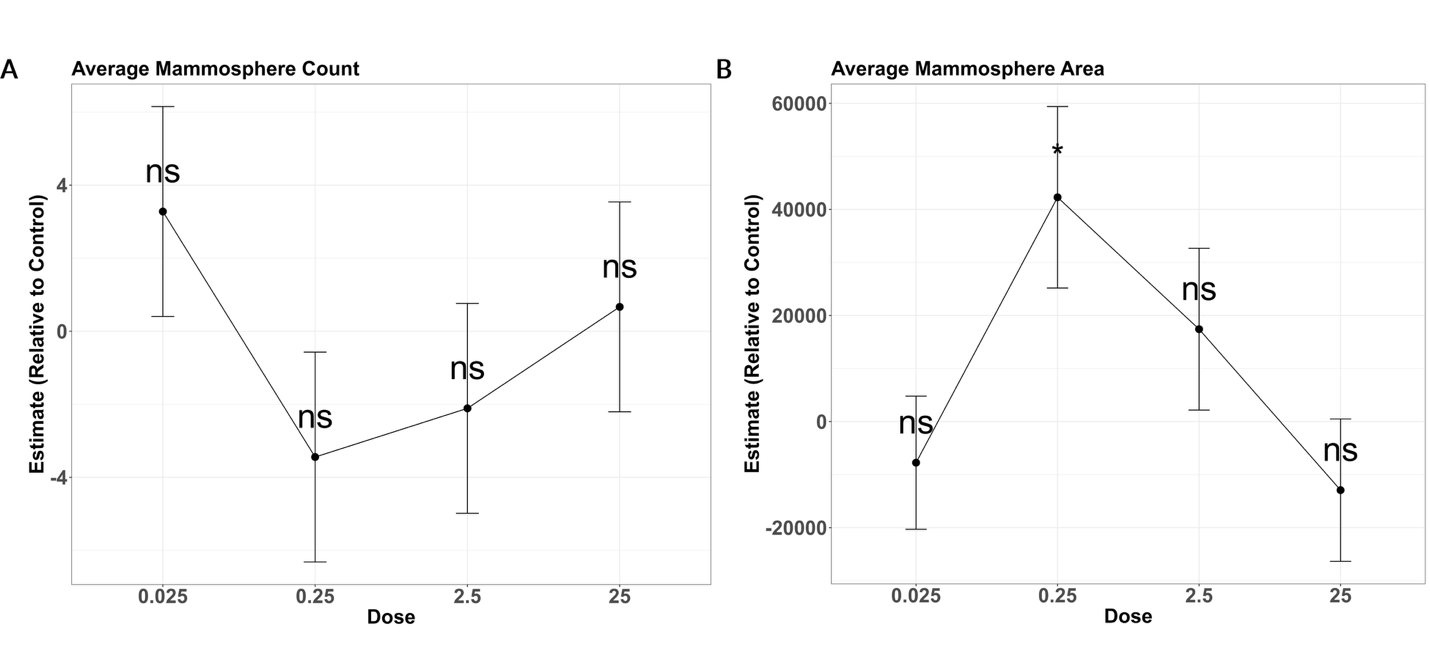

**Figure S1.** Generalized linear model results of average mammosphere count and area comparing DDE exposed mammospheres to control. A) Average mammosphere count and B) Average mammosphere area across four concentrations of DDE compared to control and across three experimental rounds. Points represent the estimated sphere number (panel A) or pixel unit area (panel B) relative to control, which is positioned at 0. Error bars represent approximate 95% confidence intervals (estimate ± 1.96 × standard error). P-values < 0.05 = *, < 0.01 = **, and < 0.001 = ***. P-values greater than 0.05 = ns.

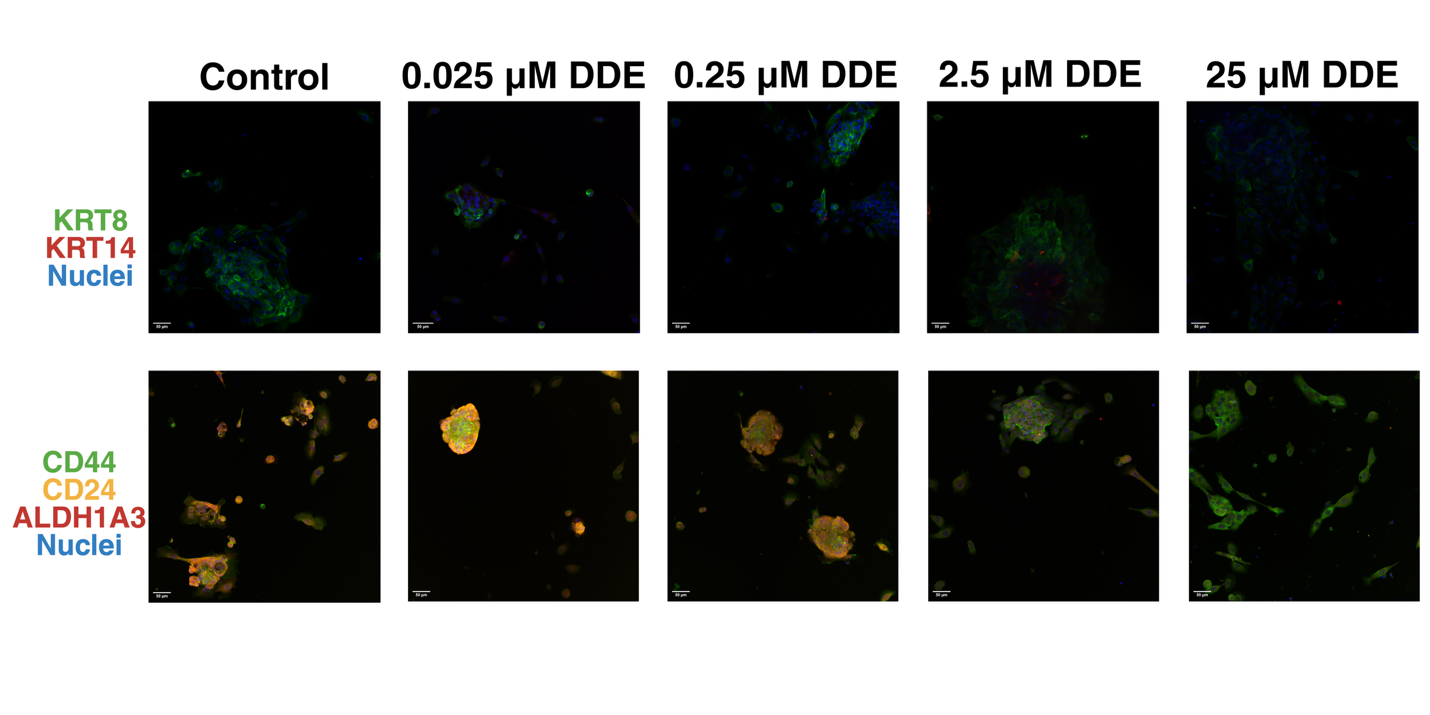

**Figure S2.** Representative images of mammospheres exposed to 0, 0.025, 0.25, 2.5, and 25 µM DDE and co-stained with KRT8 (green), KRT14 (red), and the nuclear stain Hoechst 33342 (blue) or CD44 (green), CD24 (yellow), ALDH1A3 (red), and Hoechst 33342 (blue).

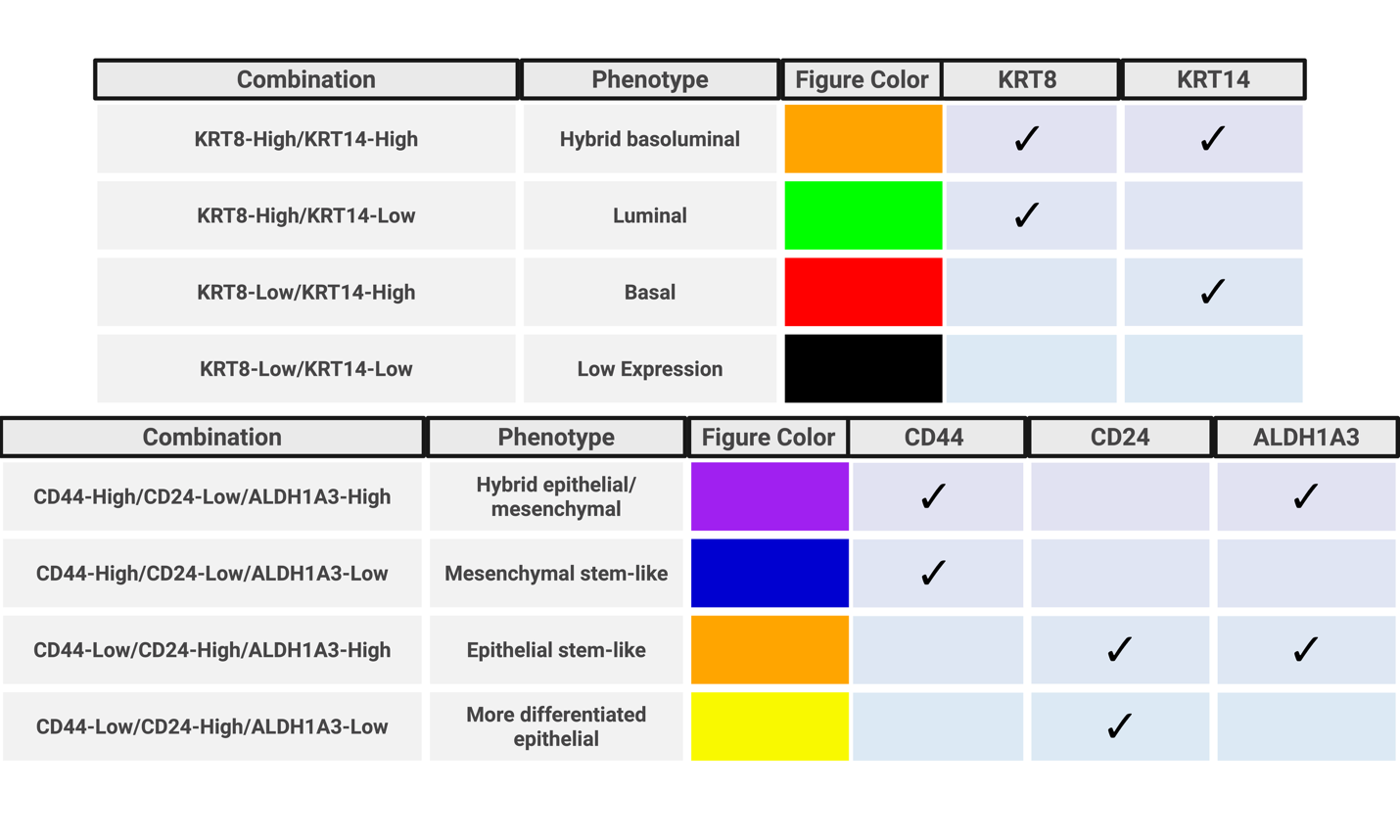

**Figure S3.** Table of cell population marker combinations, their respective phenotype, and bar color from Figure 1.

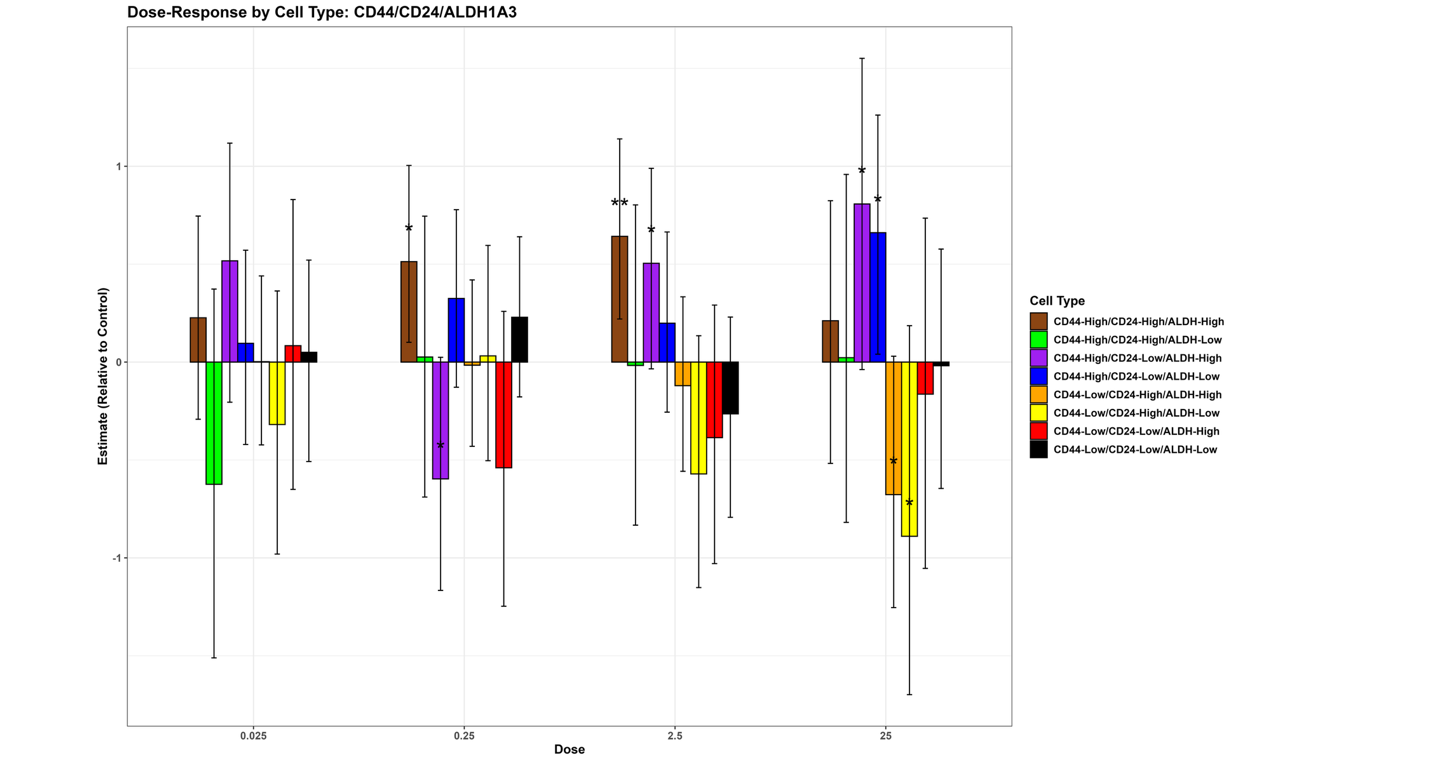

**Figure S4.** Cell-type proportions were calculated after robust normalization (median/median absolute deviation) to the control group where cells were then classified as high or low for CD44, CD24 and ALDH1A3 relative to the control median. Bars represent the posterior mean estimate (c_effect) of the cell-type composition parameter for DDE-exposed samples relative to controls, based on three experimental rounds. Error bars indicate the 95% posterior credible interval, defined by the 2.5% (c_lower) and 97.5% (c_upper) quantiles of the posterior distribution. P-values < 0.05 = *, < 0.01 = **, and < 0.001 = ***. P-values greater than 0.05 = ns.

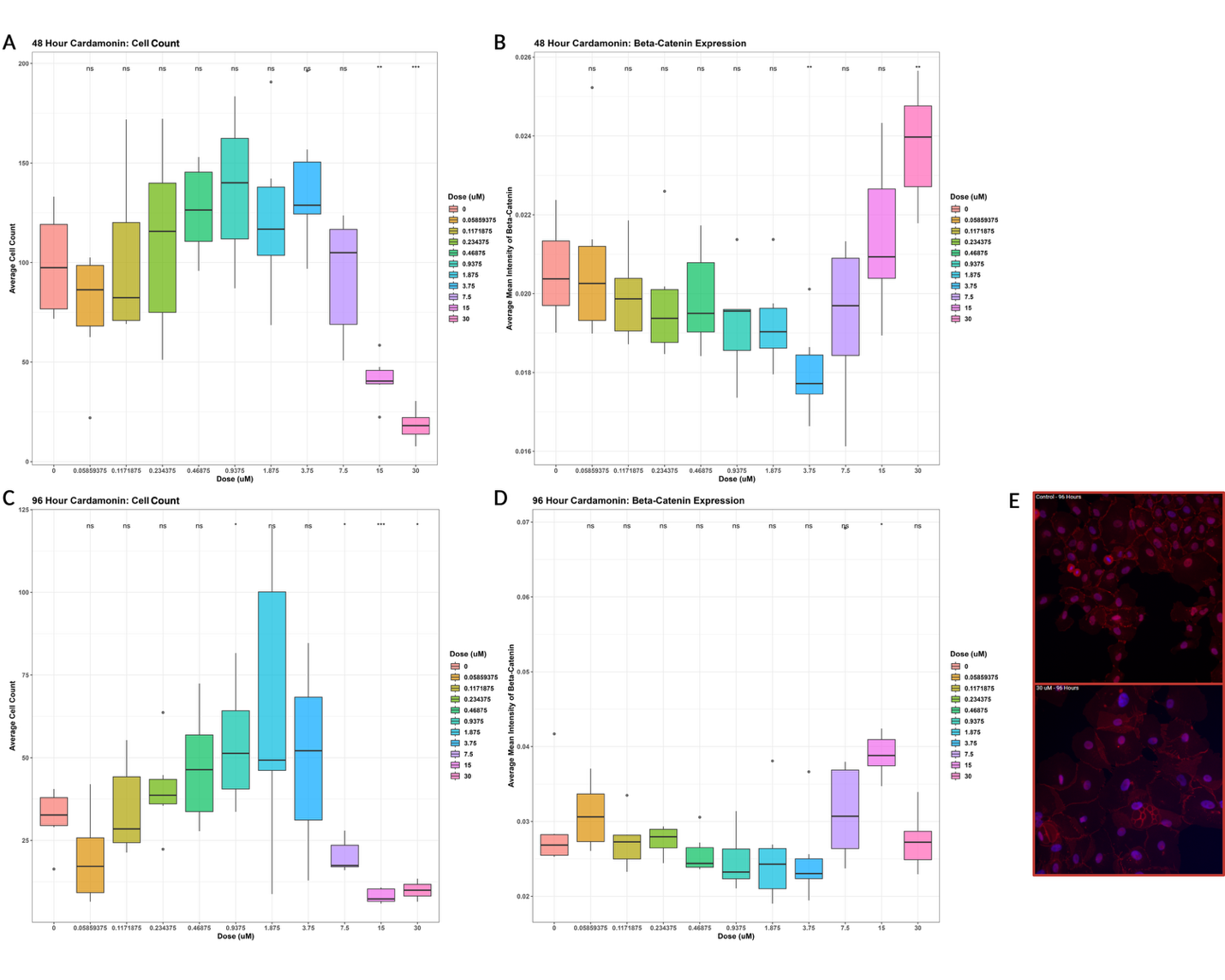

**Figure S5.** Concentration response of cardamonin exposure on MCF10A cell count (A & C) and β-catenin expression (B & D) after 48 and 96 hours. Cell counts and β-catenin expression values are averaged across six technical replicates. P-values < 0.05 = *, < 0.01 = **, and < 0.001 = ***. E) Representative images of cells stained with Hoechst 33442 and β-catenin observed after 96 hours of 30 µM cardamonin exposure compared to control. Nuclei staining with Hoechst is represented in blue and β-catenin immunostaining is represented in red.

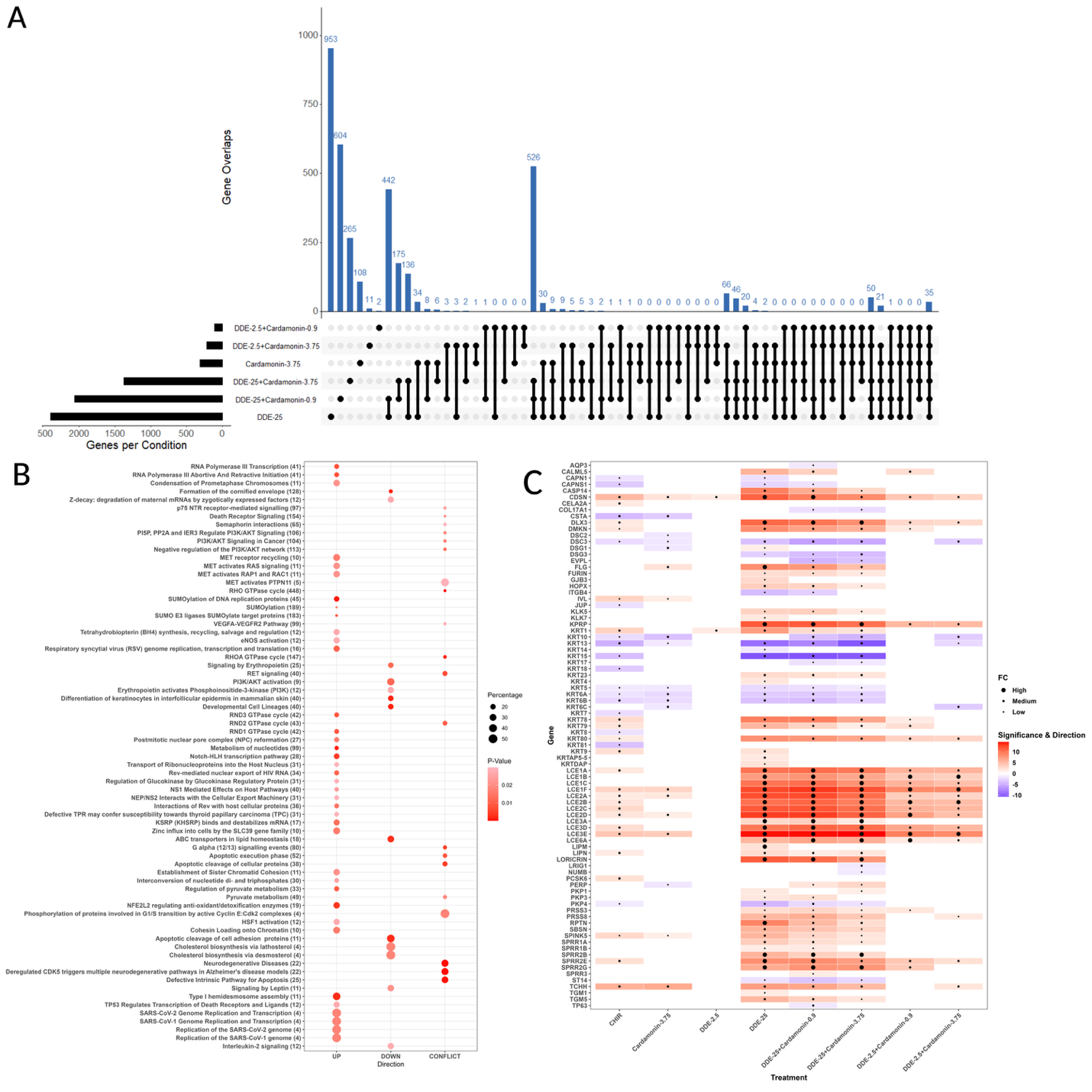

**Figure S6.** RNA-seq analysis of mammospheres exposed to DDE, cardamonin, and their mixtures. A) Upset plot of significantly differentially expressed gene overlaps, compared to control, between mammospheres exposed to 25 µM DDE (DDE-25), 3.75 µM cardamonin (Cardamonin 3.75), 25 µM DDE + 3.75 µM cardamonin (DDE-25+Cardamonin-3.75), 2.5 µM DDE + 3.75 µM cardamonin (DDE-2.5+Cardamonin-3.75), 25 µM DDE + 0.94 µM cardamonin (DDE-25+Cardamonin-0.9), and 2.5 µM DDE + 0.94 µM cardamonin (DDE-2.5+Cardamonin-0.9). The number of genes unique to each condition or group of conditions is represented by the blue bars. Each comparison is depicted below by black points and connecting lines. B) Dot plot of REACTOME pathway analysis comparing mammospheres exposed to 25 µM DDE, 25 µM DDE + 0.94 µM cardamonin, and 25 µM DDE + 3.75 µM cardamonin to assess the effects of increasing cardamonin. REACTOME pathway names are listed on the y-axis with the total number of genes in that pathway listed in parentheses. The x-axis divides the pathways that are upregulated, downregulated, or have conflicting directions of differentially expressed genes. The percentage of genes differentially expressed out of the total number of genes in the pathway is depicted by the size of the dot and only pathways with a percentage equal to or higher than 10% and FDR adjusted p-value less than 0.03 were included. FDR adjusted p-value is represented by dot color, with p-values closer to 0.03 shown as light red and p-values further from 0.03 as dark red. C) Heatmap of skin-related genes differentially expressed in each condition, including 1 µM CHIR99021, compared to control. Upregulated genes are represented by red and downregulated genes are represented by blue. Genes with greater significance are darker blue or red compared to genes with FDR adjusted p-values closer to 0.05. The log fold-change is represented by a black dot in the middle of the square, with larger logFCs being depicted as larger dots.

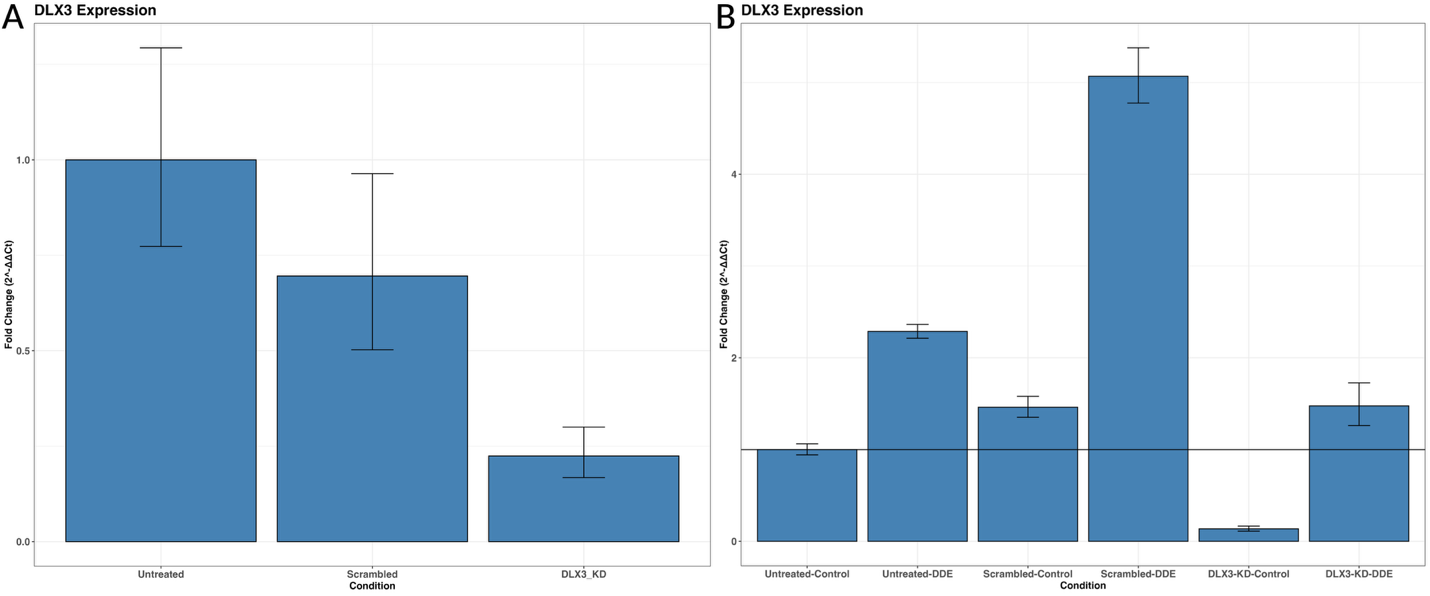

**Figure S7.** qPCR confirmation of knockdown of *DLX3*. A) Relative gene expression was calculated as 2^(-ΔΔCt) for DLX3-KD and Scrambled MCF10A cells relative to Untreated (non-transduced) MCF10A cells. Error bars were derived from the mean and standard error of the mean (SEM) for each group and calculated based on 2^(-(mean ΔΔCt +/- SEM)). B) Relative gene expression in DLX3-KD-DDE, DLX3-KD-Control, Scrambled-DDE, Scrambled-Control, and Untreated-DDE mammospheres relative to Untreated-Control mammospheres.

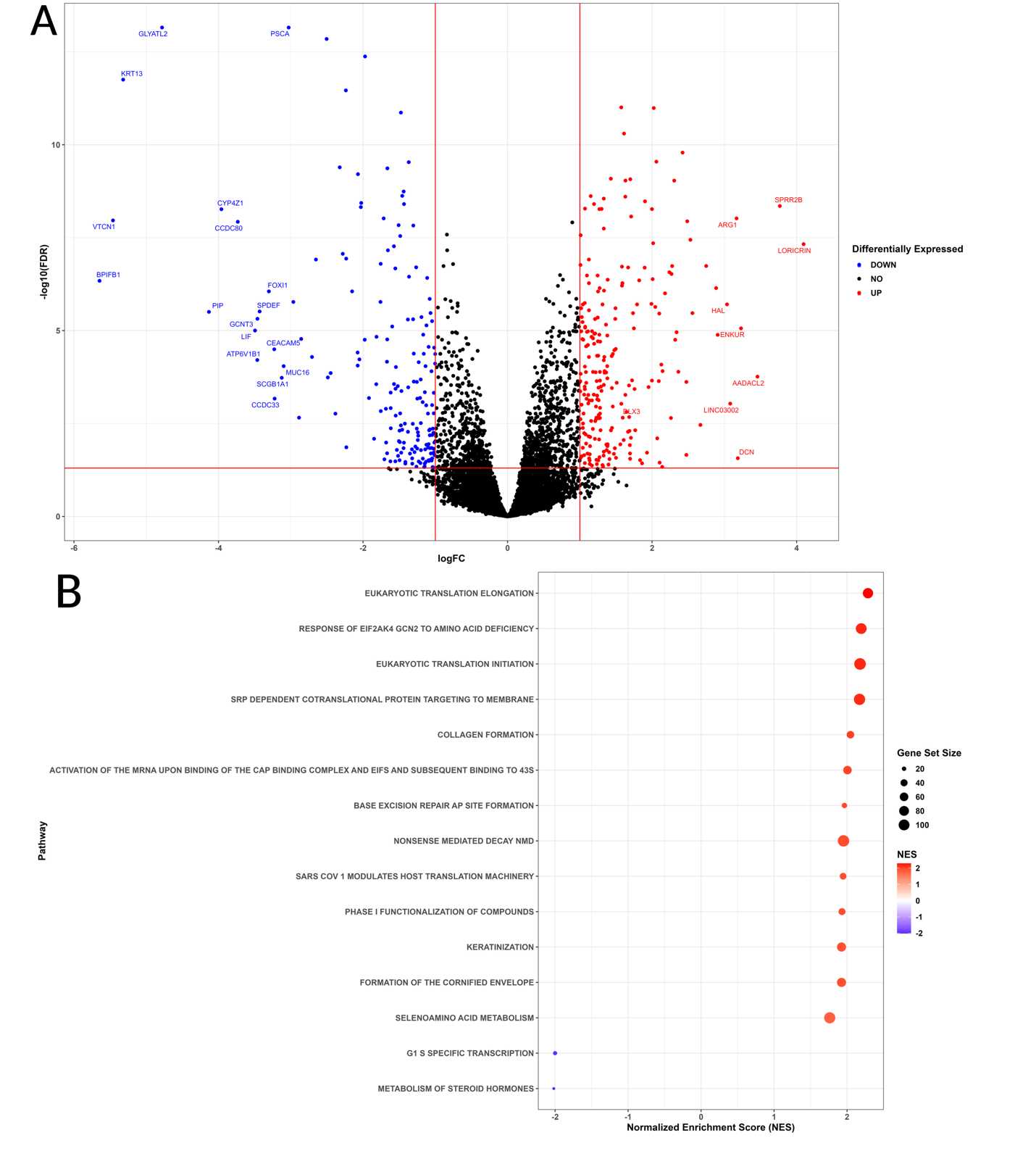

**Figure S8.** RNA-seq analysis of Scrambled-DDE (25 µM) mammospheres versus Scrambled-Control mammospheres. A) Volcano plot of differentially expressed genes. Points in red are upregulated genes compared to control and points in blue are downregulated genes compared to control. Points in black are genes that had a |log fold-change| less than 1 and/or an FDR adjusted p-value greater than 0.05. *DLX3* and all other genes with |logFCs| greater than 2 are labeled with their respective symbols. B) Dot plot of REACTOME pathway analysis comparing Scrambled-DDE mammospheres versus Scrambled-Control mammospheres. REACTOME pathway names are listed on the y-axis. The x-axis separates upregulated pathways with a positive normalized enrichment score (red) and downregulated pathways with a negative normalized enrichment score (blue). The darker shaded points represent higher absolute scores. The size of the dot represents the total number of genes in the pathway. Only pathways with an FDR adjusted p-value less than 0.05 were plotted.

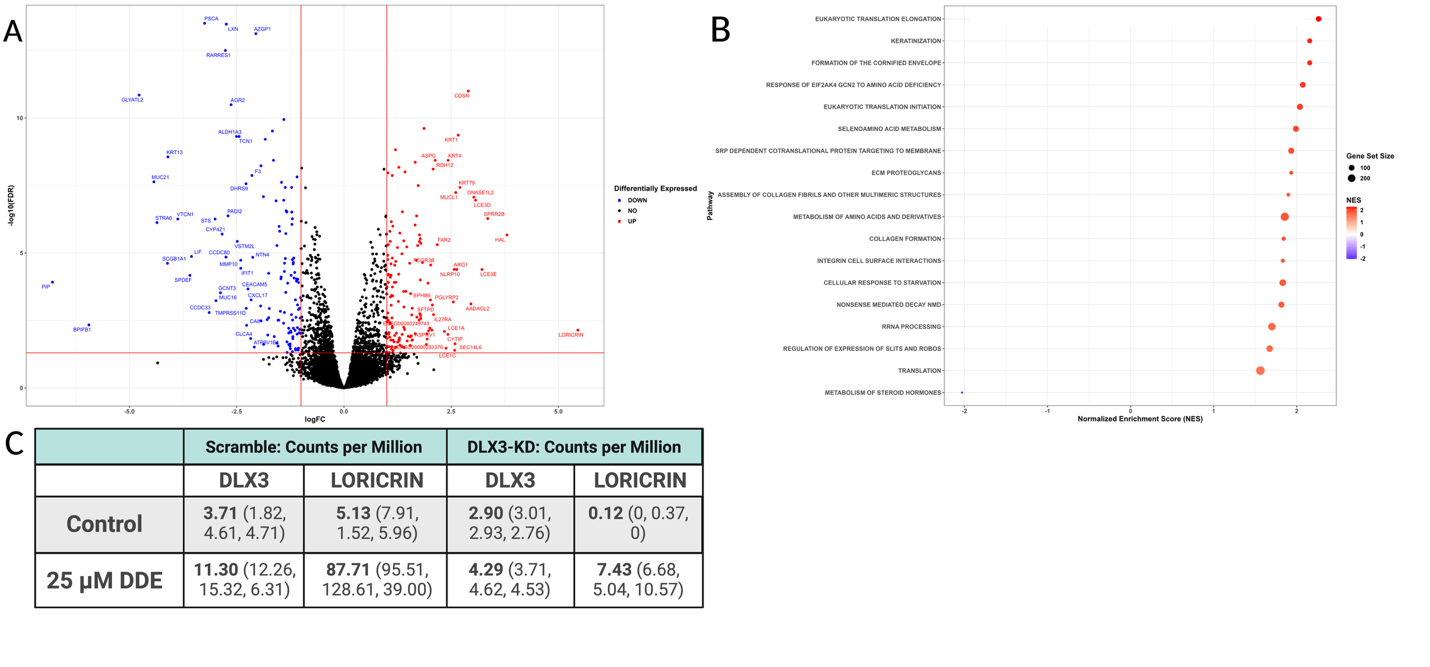

**Figure S9.** RNA-seq analysis of DLX3-KD-DDE mammospheres versus DLX3-KD-Control mammospheres. A) Volcano plot of differentially expressed genes. Points in red are upregulated genes compared to control and points in blue are downregulated genes compared to control. Points in black are genes that had a |log fold-change| less than 1 and/or an FDR adjusted p-value greater than 0.05. Genes with |logFCs| greater than 2 are labeled with their respective symbols. B) Dot plot of REACTOME pathway analysis comparing DLX3-KD-DDE mammospheres versus DLX3-KD-Control mammospheres. REACTOME pathway names are listed on the y-axis. The x-axis separates upregulated pathways with a positive normalized enrichment score (red) and downregulated pathways with a negative normalized enrichment score (blue). The darker shaded points represent higher absolute scores. The size of the dot represents the total number of genes in the pathway.  Only pathways with an FDR adjusted p-value less than 0.05 were plotted. C) Table of *DLX3* and *LORICRIN* counts per million in Scramble-Control, Scramble-DDE, DLX3-KD-Control, and DLX3-KD-DDE mammospheres. Three replicates per condition are listed with the average value in bold.

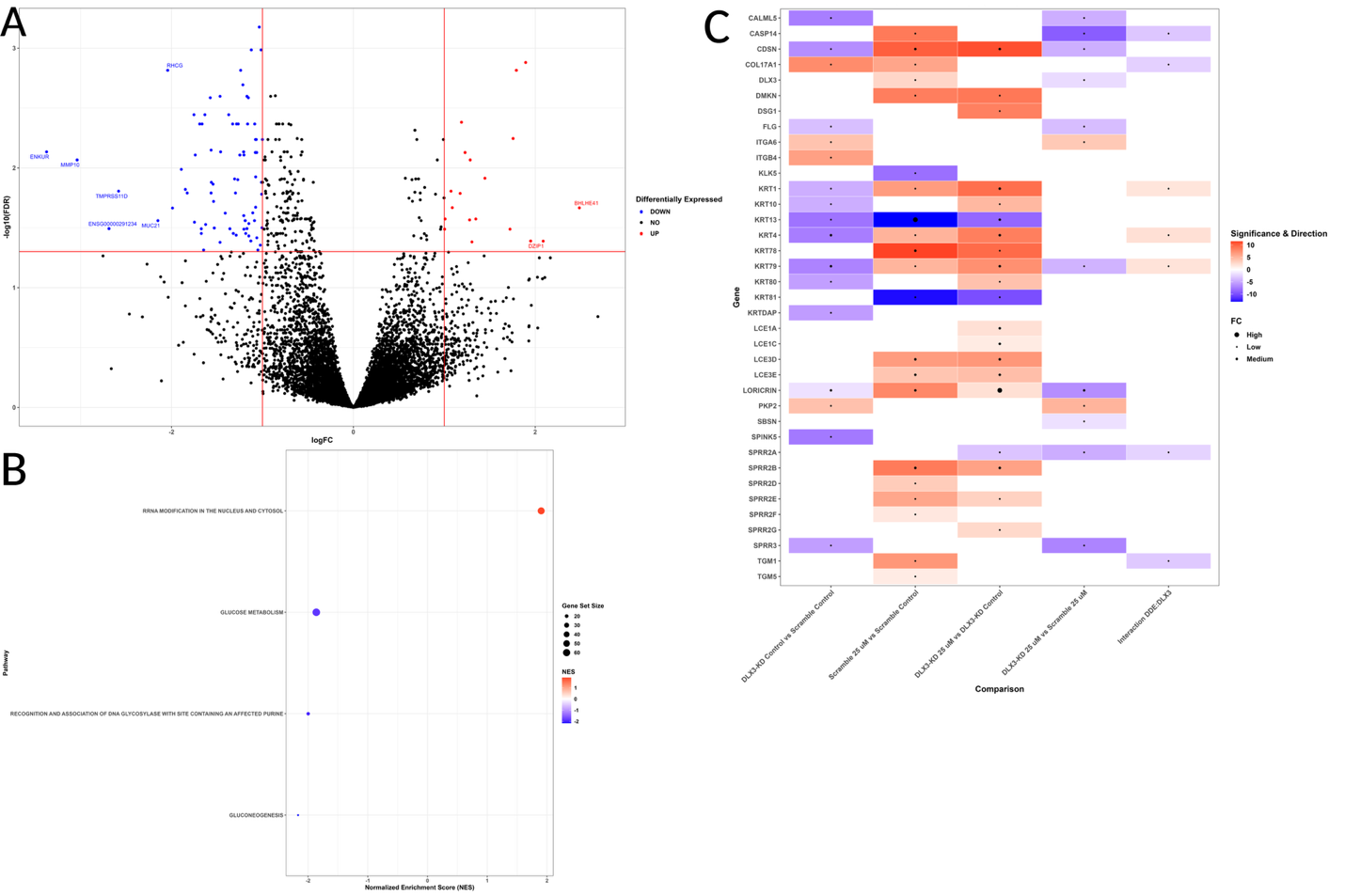

**Figure S10.** RNA-seq analysis of type-by-chemical interaction. A) Volcano plot of differentially expressed genes. Points in red are upregulated genes compared to control and points in blue are downregulated genes compared to control. Points in black are genes that had a |log fold-change| less than 1 and/or an FDR adjusted p-value greater than 0.05. Genes with |logFCs| greater than 2 are labeled with their respective symbols. B) Dot plot of REACTOME pathway analysis comparing type-by-chemical interaction. REACTOME pathway names are listed on the y-axis. The x-axis separates upregulated pathways with a positive normalized enrichment score (red) and downregulated pathways with a negative normalized enrichment score (blue). The darker shaded points represent higher absolute scores. The size of the dot represents the total number of genes in the pathway.  Only pathways with an FDR adjusted p-value less than 0.05 were plotted. C) Heatmap of skin-related genes differentially expressed in each comparison: DLX3-KD-Control vs Scrambled Control, Scrambled 25 µM vs Scrambled Control, DLX3-KD 25 µM vs DLX3-KD-Control, DLX3-KD 25 µM vs Scrambled 25 µM, and Interaction. Upregulated genes are represented by red and downregulated genes are represented by blue. Genes with greater significance are darker blue or red compared to genes with FDR adjusted p-values closer to 0.05. The log fold-change is represented by a black dot in the middle of the square, with larger logFCs being depicted as larger dots.
